# Decoding the Allosteric Paradox: A Dual Framework Integrating AI Cofolding Models with Landscape-Guided Interpretable AI Framework of Ligand-Protein Binding

**DOI:** 10.64898/2026.02.24.707829

**Authors:** Vedant Parikh, Brandon Foley, Will Gatlin, Max Ludwick, Lucas Turano, Gennady M. Verkhivker

## Abstract

Artificial intelligence (AI) has transformed prediction of protein structure and biomolecular interactions, yet modeling of allosteric regulation remains a persistent and unresolved challenge. We develop a dual explainable AI framework that systematically interrogates AI Co-Folding models AlphaFold3, Protenix, Boltz-2, Chai-1, and DynamicBind on rigorously stratified datasets of orthosteric and allosteric ligand-protein complexes. While all AI models excel in accurate modeling of orthosteric ligand binding, a universal and architecture-independent collapse emerges in prediction of allosteric complexes. The biophysical logic for this dichotomy is unveiled through physics-based lens of the energy landscape theory and local frustration analysis. Orthosteric binding creates dominant energetic funnels via ligand-induced minimal frustration quenching, while allosteric sites preserve neutral frustration landscapes in both apo and holo protein states. The findings show that conformational heterogeneity and evolutionary plasticity encoded in allosteric binding landscapes may conceal the recurrent recognition patterns AI models are trained to detect. By linking prediction outcomes to frustration landscapes, this study recasts AI shortcomings in allosteric ligand binding as diagnostic indicators of fundamental biophysical constraints, establishing a physics-informed framework that turns the allosteric blind spot into mechanistic insight for next-generation landscape-aware predictive tools.

## Introduction

AI has transformed structural biology, enabling atomic-resolution modeling of biomolecular systems and accelerating therapeutic discovery.^1–5^ This revolution began with AlphaFold2 near-experimental protein structure prediction^1,2^, extending to multi-chain complexes^6^, sequence-based protein language models (PLMs) ESMFold^7,8^ and RoseTTAFold2^9,10^, and generative design frameworks.^11^ Despite these advances, reliably predicting functional allosteric sites remains a critical limitation.^12^ Current AI models excel at recovering thermodynamically stable ground states of proteins but systematically fail to capture conformational ensembles and hidden allosteric states.^13–17^ Efforts to overcome this bias through MSA subsampling^13^, in silico mutagenesis^14^, or other sampling strategies^18–25^ have seen only partial success, constrained by training data and evolutionary priors. Concurrently, biomolecular binding prediction approaches have been shifting from purely physics-based molecular docking tools to end-to-end AI co-folding strategies. Diffusion-based architectures now jointly model protein folding and ligand placement. GeoDiff pioneered geometric denoising,^26^ TorsionDiff constrained sampling to rotatable bonds,^27^ DiffDock achieved near-experimental ligand pose accuracy in modeling of ligand-protein complexes^28^ and is further refined by implemented in its successor DiffDock-L conditional diffusion architecture.^29^ AlphaFold3 (AF3) unified this paradigm by initiating prediction from an atomic “cloud” to synchronize folding and docking across biomolecules.^30^ On the PoseBusters benchmark dataset^31^, AF3 achieved 76–81% success in recovering native ligand poses within ∼2.0 Å RMSD, substantially outperforming classical docking tools such as AutoDock Vina^32^ (52–60%) and DiffDock (∼38%)^33^. Protenix, an open-source reproduction with architectural optimizations, employs unified tokenization of polymers and small molecules^34^ and attains comparable or slightly higher success (∼80%) on PoseBusters V2^33^ . Boltz-2 extends all-atom co-folding with affinity and binder-classification heads trained on experimental and molecular dynamics (MD) simulations-derived data^35^, achieving ∼75–81% success in blind docking^33^ and leading the CASP16 Affinity Challenge.^36,37^ Chai-1 integrates paired attention with protein language model (PLM) embeddings to enable accurate predictions from sparse evolutionary data while supporting flexible loop and ligand modeling via atomic diffusion.^38^ DynamicBind advances beyond static co-folding by explicitly modeling ligand-induced protein rearrangements using an SE(3)-equivariant diffusion framework, enabling cryptic pocket formation and large-scale conformational morphing.^39^ EquiBind approach^40^ and related approach DiffSBDD^41^ use equivariant graph neural networks (GNNs) to predict ligand binding poses and pocket locations directly from apo protein structures. TANKBind approach segments proteins into functional blocks and predicts their interactions with the ligand using a trigonometry-aware architecture.^42^ Co-folding and binding AI models can now integrate protein and ligand co-design. RFdiffusion model^11^ and deep learning (DL) protein sequence design method ProteinMPNN^43^ use a protein backbone diffusion model and enable de novo design of protein binders, and small-molecule scaffolds which contain a wide range of ligand binding pockets. A multimodal foundation model ESM3 integrates sequences, structures, and functional annotations into a single architecture capable of zero-shot ligand generation.^44^ Chroma, developed by Generate Biomedicines, samples novel protein structures conditioned on desired properties.^45^ Parallel to co-folding advances, computational predictions of druggable binding sites expanded into a broad methodological ecosystem that encompasses geometric methods (Fpocket^46^, SiteHound^47^ CurPocket^48^, and MetaPocket 2.0^49^), machine learning (ML) tools such as P2Rank^50,51^ and ISMBLab-LIG^52^ that uses 3D probability density maps of interacting atoms to identify binding sites, DL architectures (DeepSite^53^, FRSite^54^, BiteNet^56^), and a spectrum of PLM-based approaches.^58–64^ Complementary physics-based approaches AlloPred^65^, Allosite^66^, AllositePro^67^, and DeepAllo^68,69^ underscored the need to embed biophysical principles within DL prediction frameworks. Despite high performance on ligand targeting orthosteric sites, all methods exhibit a pronounced and consistent drop in accuracy for allosteric binding sites and transient pockets that only form upon ligand binding.^70–72^

The central hypothesis put forward in the current study is that the emerging limitations in AI modeling of allosteric ligand binding may reflect not only algorithmic shortcomings of current AI co-folding architectures but rather represent a more fundamental biophysical mismatch between orthosteric and allosteric binding landscapes. According to this conjecture, orthosteric sites feature strong evolutionary conservation, geometric recurrence, and pronounced energetic stabilization which are signals perfectly aligned with inductive biases of AI models. Allosteric sites, by contrast, often reside in evolutionarily permissive, conformationally adaptive regions. Their functional versatility arises from the absence of dominant structural or energetic patterns which may render them to be often invisible to pattern-recognition AI models trained on recurrent motifs. These distinctions necessitate a shift to explainable biophysical AI frameworks grounded in the protein energy landscapes. A particularly informative biophysical lens is offered by local frustration analysis.^73–82^ Quantifying competing interactions within folded structures, local frustration analysis quantifies the energetic optimality of native interactions by comparing them to decoy ensembles distinguishing minimally frustrated regions optimized for stability from neutral and highly frustrated patches that are associated with conformational plasticity and functional adaptability.^73–78^ Integrating this analysis with predictive benchmarking and systematic interrogation of AI co-folding models in prediction of orthosteric and allosteric ligand binding transforms AI prediction metrics into reporters of the underlying biophysical grammar. Our analysis confirms a significant and consistent performance gap identifying the allosteric blind spot as a universal property of current architectures. We demonstrate that this divide is intrinsically linked to the distinct landscape hierarchy where orthosteric ligand binding regions are enriched in minimally frustrated, evolutionarily constrained residues, whereas allosteric ligand binding navigates through conformationally diverse and evolutionary promiscuous neutrally frustrated regions. This study demonstrates that dichotomy in prediction of orthosteric and allosteric ligand binding arises from a principled mismatch between model inductive memorization biases of AI models and the energetic logic of allosteric regulation, therefore transforming AI uncertainties into diagnostic indicators of the elusive biophysical code underlying allosteric mechanisms.

## Results

### A Dual Framework: AI Benchmarking and Explainable Energy Landscape Profiling

To systematically interrogate performance of leading AI co-folding models in prediction of orthosteric and allosteric ligand binding, we developed a dual explainable AI framework designed to distinguish between limitations imposed by training data and those imposed by the physical organization of the binding energy landscapes (Figure 1). We systematically evaluate five state-of-the-art AI co-folding models AF3^30^, Protenix^34^, Boltz-2^35^, Chai-1^38^, and DynamicBind^39^ on a rigorously curated dual benchmark of orthosteric and allosteric ligand-protein complexes obtained from Major Drug Targets (MDT) dataset^39^, rigorously curated set of allosteric ligand complexes from the Kinase Conformation Resource (KinCoRe), ^83,84^ a data set of competitive and allosteric protein kinase inhibitors confirmed by X-ray crystallography^85^ as well as validated datasets of protein-ligand orthosteric competitive complexes (PLOC) and protein-ligand allosteric (PLA) complexes.^86^

**Figure 1.**
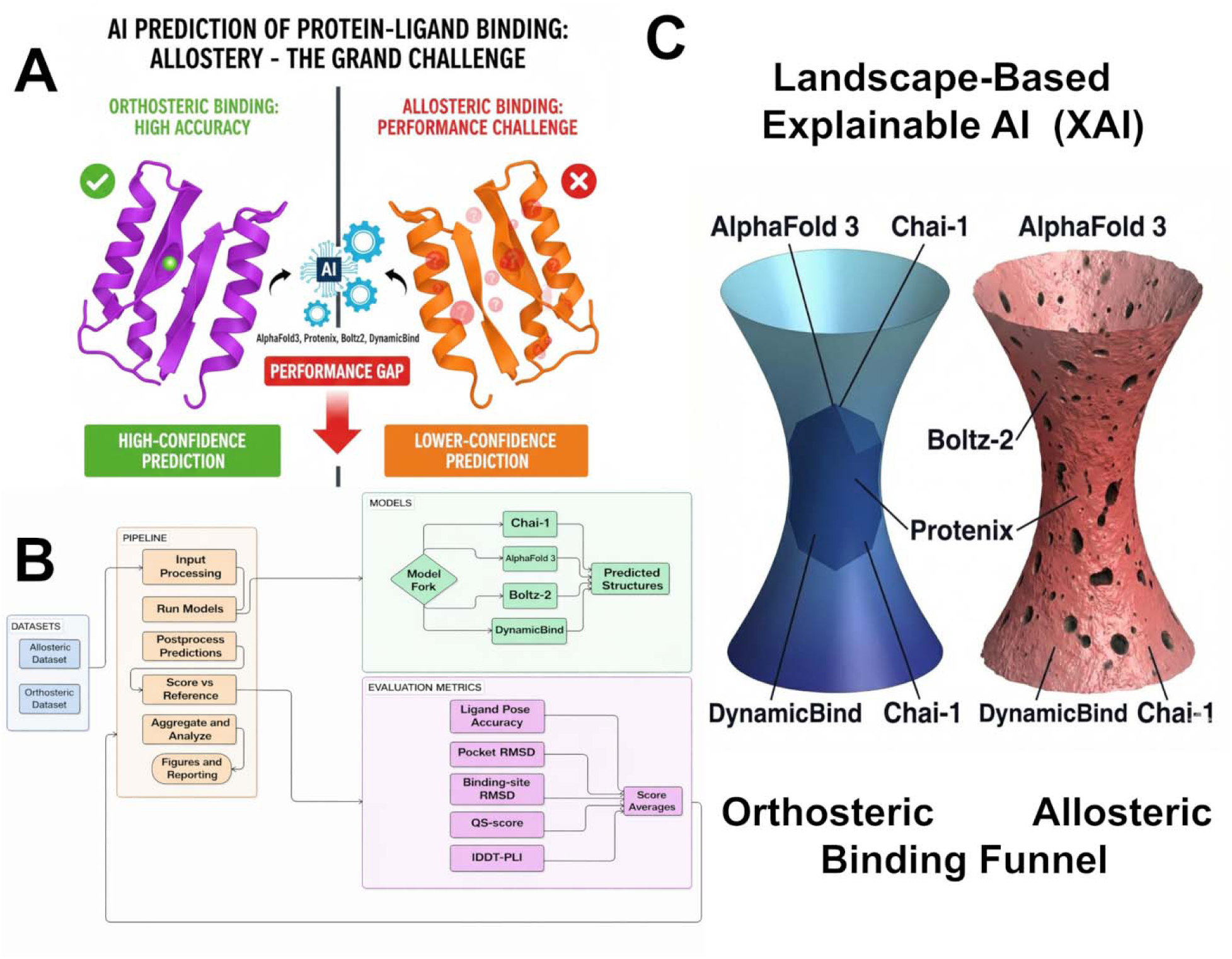
Dual explainable-AI framework for benchmarking ligand–protein binding predictions across orthosteric and allosteric sites. (A) Schematic of the explainable AI framework addressing the dichotomy in prediction of orthosteric and allosteric ligand binding. (B) Benchmarking pipeline: input processing of curated datasets, execution of five AI co-folding models (AF3, Protenix, Boltz-2, Chai-1, DynamicBind), postprocessing of predictions, scoring against reference structures, and aggregate analysis. (C) Landscape-based interpretation: orthosteric binding is governed by a single steep energetic funnel, guiding all models toward a unique high-confidence solution. Allosteric binding occurs on a flat landscape with multiple shallow minima, leading different models to predict distinct binding sites and poses.

The benchmarking component systematically evaluates performance on rigorously stratified orthosteric and allosteric datasets, assessing two interconnected tasks that for allosteric ligand binding present a compounded challenge : identification of the ligand binding pocket and structural prediction of the ligand binding pose within the pocket. The interpretability layer based on conformational ad mutational frustration analyses provides a quantitative readout of the energetic and evolutionary organization of the ligand binding enabling direct testing of the hypothesis that robustness and accuracy of allosteric ligand predictions can be dictated not by model architecture but by the topological organization of the underlying energy landscape. We demonstrate that orthosteric binding sites undergo frustration quenching upon ligand binding, transitioning from elevated apo-state frustration to minimal frustration upon ligand binding and creating steep landscape funnels serving as unambiguous signals for AI convergence. In contrast, allosteric binding can preserve neutral frustration in both apo and holo states that obscures structural and evolutionary patterns that AI models are trained to detect thus underlying the biophysical logic behind model-independent performance gap. By linking quantitative predictions of AI models to the landscape-based explainable AI framework, this study recasts allosteric prediction bind spots not only as algorithmic shortcomings but rather as diagnostic indicators of biophysical allosteric grammar.

### Orthosteric Binding Exhibits Unimodal Convergence to Near-Experimental Accuracy Across Diverse Architectures

We first sought to establish a high-fidelity baseline for canonical orthosteric binding against which allosteric performance could be compared. To this end, we assembled a comprehensive orthosteric reference combining the MDT dataset (412 complexes from kinases, GPCRs, nuclear receptors, and ion channels selected for structural challenge requiring pocket RMSD greater than 2.0 Å between apo and holo structures) together with the PLOC subset adding 1,863 experimentally determined structures with 2,325 binding pockets and 1,647 unique ligands spanning diverse families, all binding competitively to primary, evolutionarily conserved orthosteric sites.^86^ The combined orthosteric reference totals 2,275 complexes, capturing the diversity of evolutionarily optimized, geometrically well-defined binding sites. Across all five AI models evaluated on this comprehensive reference, orthosteric ligand pose predictions converged tightly around native geometries (Figure 2A). AF3, Protenix, and Chai-1 achieved mean ligand pose RMSD of 2.3–2.8 Å; Boltz-2 and DynamicBind reached mean ligand pose RMSD of 3.8–4.1 Å (Figure 2A). Pocket RMSD analysis revealed complementary structural fidelity (Figure 2B). DynamicBind achieved the lowest mean pocket RMSD of approximately 1.0 Å while AF3 and Protenix maintained mean pocket RMSD were below 2.0 Å (Figure 2B). Beyond geometric accuracy, orthosteric predictions achieved near-perfect recovery of native contact topology, with QS-scores exceeding 0.90 across all models (Figure 2C) further reinforcing the robust encoding of orthosteric ligand-protein binding pockets and interaction patterns in training data. At the stringent threshold of 2.0 Å for ligand pose RMSD, both AF3 and Protenix models exceeded 80% success rates, a benchmark reflected in the success rate matrix (Figure 2D), which shows consistently high performance accuracy of orthosteric binding predictions across all models.

**Figure 2.**
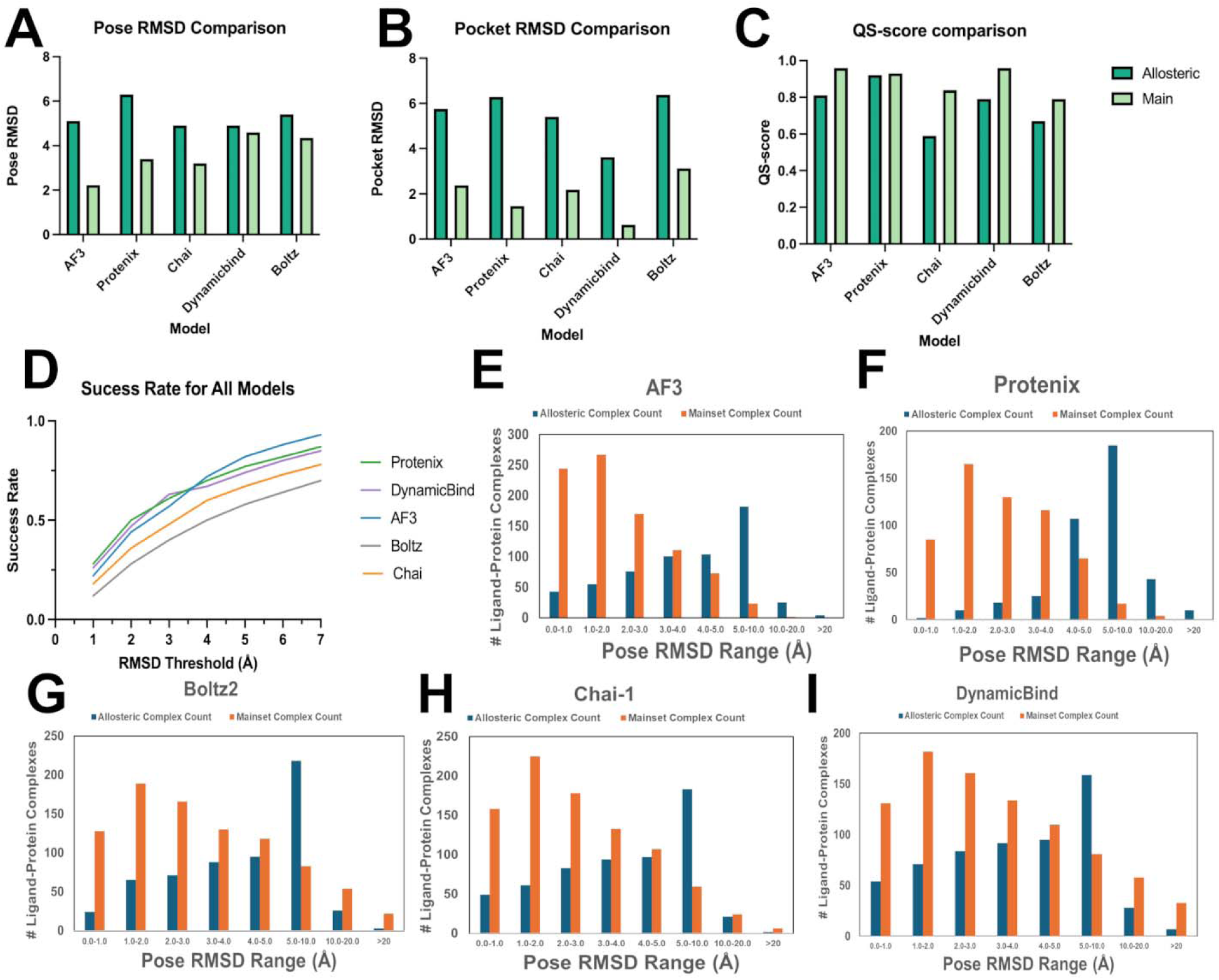
Architecture-independent performance collapse for allosteric binding reveals compounded failure in pocket localization and pose prediction. (A) Mean ligand pose RMSD. Orthosteric complexes are predicted with near-experimental accuracy across all five AI models (RMSD = 2.3–4.1 Å; light green bars), whereas allosteric complexes exhibit a systematic and architecture independent degradation, with mean RMSD increasing to 5.2–6.8 Å (dark green bars). (B) Mean pocket RMSD. Structural fidelity of the binding pocket mirrors the pose degradation. Orthosteric pocket RMSD remains low (1.4–3.0 Å), whereas allosteric pocket RMSD nearly doubles (3.5–6.3 Å). (C) QS-score comparison. Orthosteric complexes achieve near-perfect recovery of native contact topology (QS > 0.90), while allosteric complexes retain moderately high QS-scores (0.70–0.85) revealing a systematic decoupling between interaction topology and precise pose convergence. (D) Success rate matrix. Fraction of predictions achieving ligand pose RMSD < 2.0 Å across models and datasets, illustrating the dramatic drop in success rates from orthosteric to allosteric targets. (E–I) Granular RMSD distributions. Binned ligand pose RMSD histograms for AF3 (E), Protenix (F), Boltz-2 (G), Chai-1 (H), and DynamicBind (I). Orthosteric predictions (orange) exhibit sharp unimodal peaks with >70% of predictions within 0–2 Å, consistent with convergence guided by strong energetic funnels. In contrast, allosteric predictions (blue) collapse into broad, flat distributions with dominant mass shifted to 5–20 Å.

The most striking feature emerges from granular distribution analysis (Figures 2E–G). Across all models, more than 70% of predictions cluster within ligand pose RMSD ∼ 0–2 Å, forming a sharp unimodal distribution centered on the native pose. This is the signature of a well-defined energetic funnel as the landscape geometry forces sampling trajectories toward a single dominant minimum, enabling different architectural approaches to arrive at the same experimentally validated solution.

### Allosteric Binding Induces Architecture-Independent Performance Collapse

The same five AI models were evaluated on the assembled allosteric dataset integrating kinase-specific complexes from the KinCoRe database ^83,84^ (217 Type III/IV allosteric inhibitors), a curated set of high-confidence allosteric kinase inhibitors (136 complexes)^85^, and the PLA subset comprising 1,613 non-kinase structures across diverse protein families.^86^ The results revealed a systematic and profound collapse in predictive performance. Mean ligand pose RMSD increased to 5.2–6.8 Å across all architectures, effectively doubling the orthosteric error range of 2.3–4.1 Å (Figure 2A). This degradation is not model-specific identifying the allosteric blind spot as a universal phenomenon rooted in the nature of allosteric binding landscape rather than in algorithmic design.

Importantly, this collapse affects both components of the dual prediction task. Mean pocket RMSD nearly doubled, increasing from 1.4–3.0 Å for orthosteric sites to 3.5–6.3 Å for allosteric sites (Figure 2B). Protenix shows the most severe pocket degradation, with mean pocket RMSD rising from 1.5 Å to 6.2 Å, while even DynamicBind, which is the strongest performer for pocket localization, exhibits a substantial increase from RMSD 1.0 Å to 3.8 Å (Figure 2B). AI models are often unable to accurately localize regulatory pockets, and even when pockets are approximately identified, AI predictions of allosteric ligand binding poses may deviate significantly from experimental solutions -0 a compounded failure that the QS-score analysis further illuminates (Figure 2C). Indeed, while orthosteric complexes achieve near-perfect contact recovery with QS-scores exceeding 0.90, allosteric QS-scores remain moderately high at 0.70–0.85 despite catastrophic geometric error, revealing that models may identify the correct interaction partners but cannot accurately resolve their spatial arrangement into a unique binding geometry. This indicates that even when the proper interaction partners are recognized for allosteric ligands, the energetic guidance is often insufficient to enforce geometric convergence. Success rates for allosteric ligand binding at the stringent 2.0 Å ligand pose RMSD threshold collapse to 25–35% across AI models (Figure 2D), and the tightly unimodal orthosteric distributions transform into broad, flat profiles extending towards RMSD > 10 Å (Figures 2E–G). AF3 predictions skew strongly toward the RMSD - 5–10 Å bin with a pronounced long tail beyond 20 Å. Chai-1 displays a modest peak at RMSD= 2–3 Å followed by rapid decay, consistent with the inability of sequence-based priors to compensate for missing coevolutionary signals at evolutionarily permissive allosteric sites. Boltz-2 exhibits the most severe degradation, with predictions concentrated in the RMSD = 5–10 Å range. DynamicBind retains a small peak at RMSD= 2–3 Å for allosteric ligands, reflecting its emphasis on induced-fit modeling, yet still fails for the majority of complexes. Model-specific patterns in pose degradation are also evident (Figure 2A). AF3 and Protenix, the top performers on orthosteric sites, exhibit the largest absolute degradation, with mean ligand pose RMSD increasing by approximately 4.1 Å (Figure 2A). DynamicBind shows comparative resilience with mean ligand pose RMSD ∼ 4.5 Å yet remains far from experimental accuracy, underscoring that explicit flexibility modeling may be insufficient to overcome fundamental biophysical constraints. Intriguingly, architectural features of AI models optimized for extracting maximal signal from high-quality orthosteric input could become liabilities when confronted with energetic degeneracy of allosteric binding.

Together, these results demonstrate that the observed performance gap of AI models on allosteric ligand binding arises from a compounded challenge of simultaneous mislocalization of binding pockets and lack of geometric convergence, revealing the universal “allosteric blind spot” across diverse AI architectures that may reflect fundamental differences in the biophysical organization of binding sites rather than model-specific limitations.

Figure 3 further illustrates the fundamental contrast between orthosteric and allosteric ligand binding predictions across all evaluated AI models. For all five models, orthosteric ligand pose RMSD (Figure 3A-E), pocket RMSD (Figure 3F-J), and QS-score distributions (Figure 3K-O) exhibit narrow, symmetric distributions centered below RMSD ∼3.0 Å, indicating high prediction precision and unimodal convergence toward the native pose. Narrow interquartile ranges and the absence of extended tails indicate robust geometric convergence and demonstrate that orthosteric binding constitutes a well-posed prediction problem for AI models. In contrast, allosteric predictions display broad, flattened distributions extending beyond RMSD = 10 Å, reflecting systematic loss of geometric accuracy and the emergence of multiple distinct, non-native solutions. AF3 and Protenix show the strongest broadening for allosteric sites, whereas DynamicBind exhibits the most compressed allosteric distribution (Figure 3).

**Figure 3.**
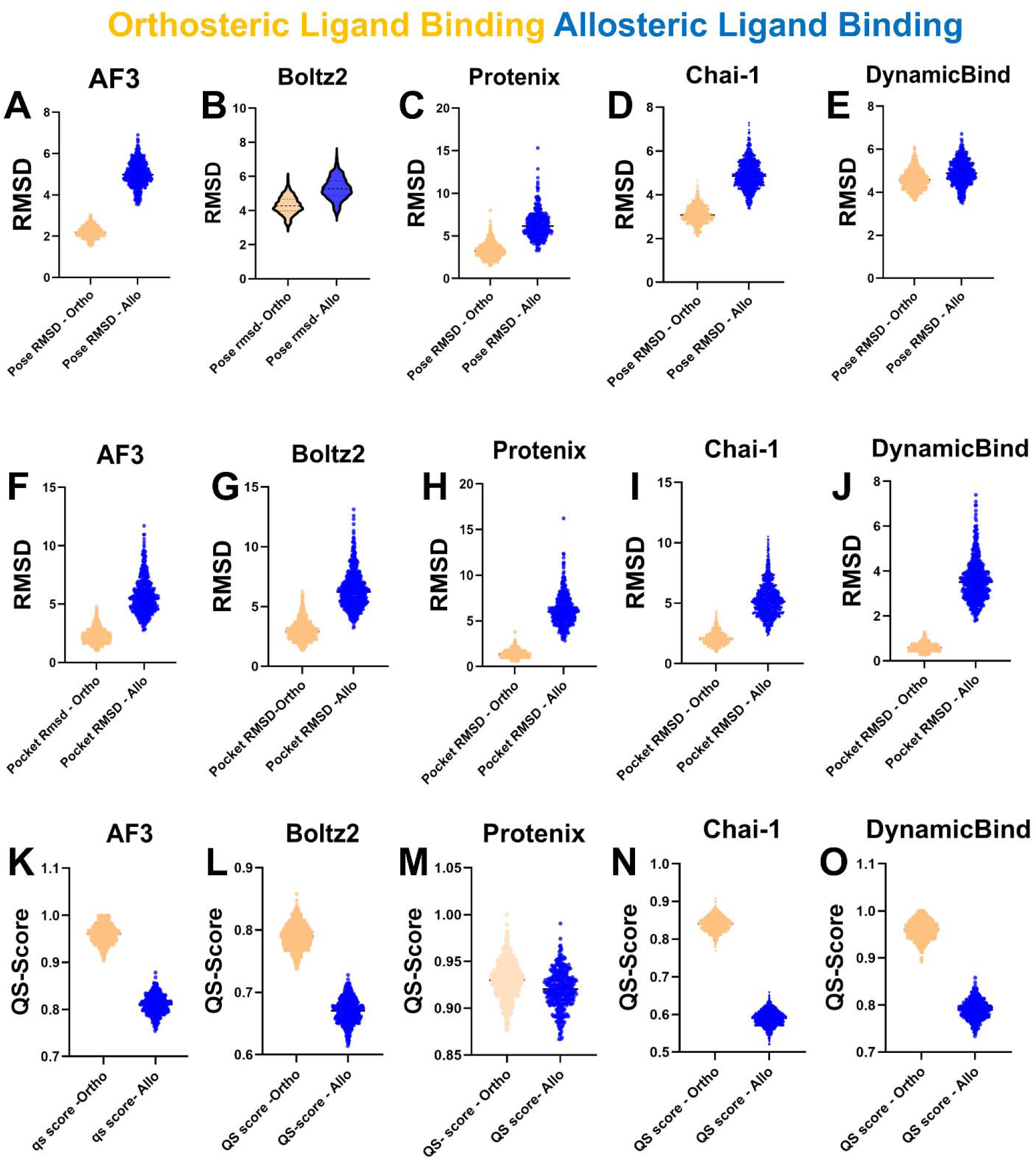
Distributional comparison of AI model performance for orthosteric versus allosteric ligand binding across pose accuracy, pocket localization, and contact topology. Violin plots summarize prediction performance for five AI co-folding models (AF3, Protenix, Boltz-2, Chai-1, and DynamicBind), contrasting orthosteric complexes (orange) with allosteric complexes (blue). (A–E) Ligand pose RMSD distributions. For all five models, orthosteric predictions (orange) exhibit narrow, symmetric distributions centered below RMSD ∼3.0 Å, indicating high geometric precision and unimodal convergence toward the native pose. In contrast, allosteric predictions (blue) display broad, flattened distributions extending beyond RMSD = 10 Å, reflecting systematic loss of geometric accuracy and the emergence of multiple distinct, non-native solutions. AF3 and Protenix show the strongest broadening for allosteric sites, whereas DynamicBind exhibits the most compressed allosteric distribution, consistent with its partial resilience to induced-fit effects. (F–J) Pocket RMSD distributions. Orthosteric binding pockets (orange) are localized with higher precision, yielding tight distributions typically below ∼3.0 Å across models. Allosteric pockets (blue) show markedly wider and right-shifted distributions, confirming difficulty in correctly identifying diverse and weakly conserved allosteric binding sites. (K–O) QS-score (contact topology) distributions. Orthosteric complexes (orange) achieve near-perfect recovery of native contact networks (QS > 0.90) for all models, indicating faithful reconstruction of interaction topology. Allosteric complexes (blue) retain only moderate QS-scores (0.70–0.85) revealing a systematic decoupling between contact topology recovery and geometric precision.

This divergence is particularly pronounced for AF3 and Protenix: their sharply peaked orthosteric violins (median RMS ∼ 2.5 Å, interquartile range <1.5 Å) transform into highly diffuse allosteric distributions spanning approximately RMSD = 2–20 Å, with medians exceeding RMSD = 6 Å.

Allosteric pocket RMSD distributions exhibit a parallel expansion, with medians shifted to RMSD valu3es ∼ 3.5–5.0 Å and long tails extending beyond RMSD of 8 Å. Importantly, this broadening is heterogeneous: while some allosteric pockets are localized with near-orthosteric precision, others are completely missed. This heterogeneity reflects the architectural diversity of allosteric sites and the lack of consistent geometric or evolutionary signatures that AI models can reliably exploit for pocket identification.

Across all panels, the persistent separation between orthosteric and allosteric distributions reveals an architecture-independent “allosteric blind spot.” Orthosteric sites are governed by steep energetic funnels that enforce unimodal convergence toward a unique native pose, whereas allosteric sites reside on flatter or degenerate landscapes that permit multiple geometrically distinct solutions with comparable interaction topologies. This energetic degeneracy drives systematic underperformance across diverse AI architectures and underlies the intrinsic difficulty of allosteric ligand binding prediction. The violin distributions in Figure 3 demonstrate that prediction accuracy for orthosteric and allosteric complexes differs not only in magnitude but in statistical structure. Orthosteric predictions form tight, symmetric distributions with narrow interquartile ranges and short tails across all three metrics—ligand pose RMSD, pocket RMSD, and QS-score—indicating that binding geometry and interaction topology are simultaneously well constrained. In contrast, allosteric distributions are broad, flat, and frequently bimodal, with long tails extending to high RMSD values while QS-scores remain comparatively elevated. This pattern suggests that allosteric prediction failure is not simply a matter of increased noise or random error but can reflect a systematic divergence between interaction recovery and geometric convergence. To directly evaluate this hypothesis, we next examine whether AI models can recover correct contact topology even when geometric accuracy collapses by analyzing QS-scores independently of pose RMSD.

### Contact Topology Recovery Decouples from Geometric Precision in Allosteric Complexes

To disentangle geometric failure from interaction failure, we quantified recovery of native contact topology using QS-scores, which measure agreement between predicted and experimental interaction networks independently of absolute pose geometry. For orthosteric complexes, QS-scores exceed 0.90 across all five models, with AF3 and Protenix approaching 0.95 (Figure 2C). This near-perfect recovery demonstrates that canonical orthosteric interaction patterns are robustly encoded and tightly coupled to geometric accuracy: when models predict correct contacts, they also converge to the native pose. In sharp contrast, allosteric complexes exhibit a striking decoupling between topology and geometry. Despite ligand pose RMSD frequently exceeding 5 Å, QS-scores remain moderately high (0.70–0.85) for AF3, Protenix, and DynamicBind (Figure 2C). Allosteric QS-score distributions show only a modest downward shift relative to orthosteric values and remain relatively tight, even as geometric error becomes extreme. This indicates that models frequently identify the correct regulatory region and reconstruct qualitatively native interaction networks but fail to resolve these interactions into a unique geometric arrangement.

This decoupling is visualized systematically in the bulb scatter plots (Figures S1 and S2). For orthosteric complexes (Figure S1), predictions cluster densely in the low-RMSD (<3 Å), high-QS (>0.9) quadrant, reflecting strong coupling between interaction topology and geometry.

For allosteric complexes (Figure S2), the distribution spreads dramatically: ligand and pocket RMSDs span a wide range of 4–10 Å, while QS-scores remain elevated (0.7–0.9). This pattern is conserved across all five architectures, confirming that the decoupling is not model-specific but reflects an intrinsic property of allosteric binding. This behavior provides the central mechanistic insight of the benchmark. Allosteric binding prediction fails not because models cannot identify relevant interaction partners, but because they cannot resolve the energetic specificity required for geometric convergence. Evolutionary pressures acting on allosteric sites prioritize regulatory flexibility and ensemble redistribution rather than tight energetic optimization, yielding landscapes in which multiple geometrically distinct conformations satisfy similar interaction topologies. Based on these findings, we argue that AI models can recover the vocabulary of allosteric interactions (which residues contact the ligand) but cannot determine the syntax (their precise spatial arrangement), because several syntactically distinct solutions are energetically comparable. This interpretation is exemplified by representative structures in Figure S3. For orthosteric complexes, predicted poses align closely with experimental conformations across architectures, typically within RMSD =1–2 Å (Figure S3). For allosteric complexes, predictions diverge substantially, adopting alternative ligand orientations or misplacing the pocket entirely, even when native contacts are partially preserved.

Granular analysis further reveals that a substantial fraction of orthosteric predictions reach the sub-angstrom regime, approaching crystallographic uncertainty (Figure S4). AF3 achieves the highest proportion of ultra-precise predictions, with 72% of orthosteric complexes below 2.0 Å RMSD and a large fraction below 1.0 Å, reflecting its ability to exploit steep energetic funnels. Chai-1 shows comparable enrichment in the high-precision regime, consistent with its reliance on evolutionary priors that reinforce canonical geometries. Boltz-2 exhibits fewer sub-angstrom predictions, reflecting its emphasis on confidence calibration rather than maximal geometric refinement (Figure S4). This hierarchy in which AF3 and Chai-1 lead in the ultra-high precision regime, followed by Boltz-2 mirrors the architectural tradeoffs observed throughout this study. The sub-angstrom regime represents the upper echelon of what current AI architectures can achieve when the underlying energy landscape provides an unambiguous directional signal. Models that excel at extracting maximal signal from steep funnels achieve exceptional accuracy on orthosteric sites, but this very sensitivity becomes a liability when confronted with the promiscuous landscapes of allosteric binding. QS-score analysis reveals a fundamental dichotomy. Orthosteric sites exhibit tight coupling between contact topology and geometry, near-perfect QS-scores, unimodal RMSD distributions, and uniform performance across architectures—hallmarks of binding governed by steep energetic funnels and frustration quenching. Allosteric sites display the opposite pattern: doubled pose RMSD, broadened pocket error, and persistent topology–geometry decoupling, with QS-scores remaining moderately high despite catastrophic geometric divergence.

Taken together, the benchmarking results reveal a striking and reproducible dichotomy across metrics, datasets, and architectures. For orthosteric sites, the evidence is unequivocal: unimodal RMSD distributions centered at 1–2 Å, tight coupling between pocket localization and pose accuracy, near-complete recovery of contact topology (QS > 0.9), and uniform performance across all five models. For allosteric sites, an equally consistent but opposite regime emerges. Mean ligand pose RMSD doubles to 5–7 Å, success rates at the 2.0 Å threshold collapse to 25–35%, and narrow unimodal distributions broaden into flat or multimodal profiles extending beyond 20 Å. Pocket RMSD increases in parallel, and although contact topology recovery remains moderately high (QS = 0.70–0.85), this stands in sharp contrast to the accompanying geometric failure. Despite fundamentally different inductive biases across A models, the architecture-independent nature of this collapse indicates that the allosteric blind spot is unlikely to be algorithmic in origin. Instead, it points to intrinsic biophysical differences in the structural and evolutionary organization of orthosteric and allosteric sites.

### Energy Landscape Architecture Encodes the Physical Basis of Explainable AI: Ligand-Induced Frustration Quenching Creates Energetic Funnels for Orthosteric Ligand Binding

By overlaying AI prediction outcomes with frustration-based landscape profiles, we link model performance on orthosteric and allosteric ligand binding to the physical organization of their underlying energy landscapes, transforming benchmark accuracy into mechanistic explanation. This approach addresses a central question raised by our benchmarking results: which features of allosteric sites render them resistant to accurate prediction, and conversely, what features of orthosteric sites make them so reliably detectable? To answer this question, we employed local frustration analysis, an energy landscape–theory framework that quantifies the energetic optimality of native interactions relative to large decoy ensembles. This analysis provides complementary residue-level descriptors of (i) conformational frustration, which reports on local landscape topography relevant to pose placement, and (ii) mutational frustration, which reflects evolutionary constraint encoded in sequence and thus informs pocket detectability. The frustration analysis was explicitly designed to mirror the dual prediction task faced by AI models. Our analysis separately interrogates (i) the intrinsic energetic organization of binding sites in the apo state, which governs detectability, and (ii) the energetic reorganization induced by ligand binding, which governs geometric convergence. By comparing apo and holo states across orthosteric and allosteric complexes, we distinguish features intrinsic to binding sites from those that emerge only upon ligand engagement.

We first examined conformational frustration across entire protein folds to establish a global baseline. Orthosteric and allosteric complexes displayed nearly identical global frustration profiles in both apo and holo states (Figure 4A–B for orthosteric; Figure 4E–F for allosteric).

**Figure 4.**
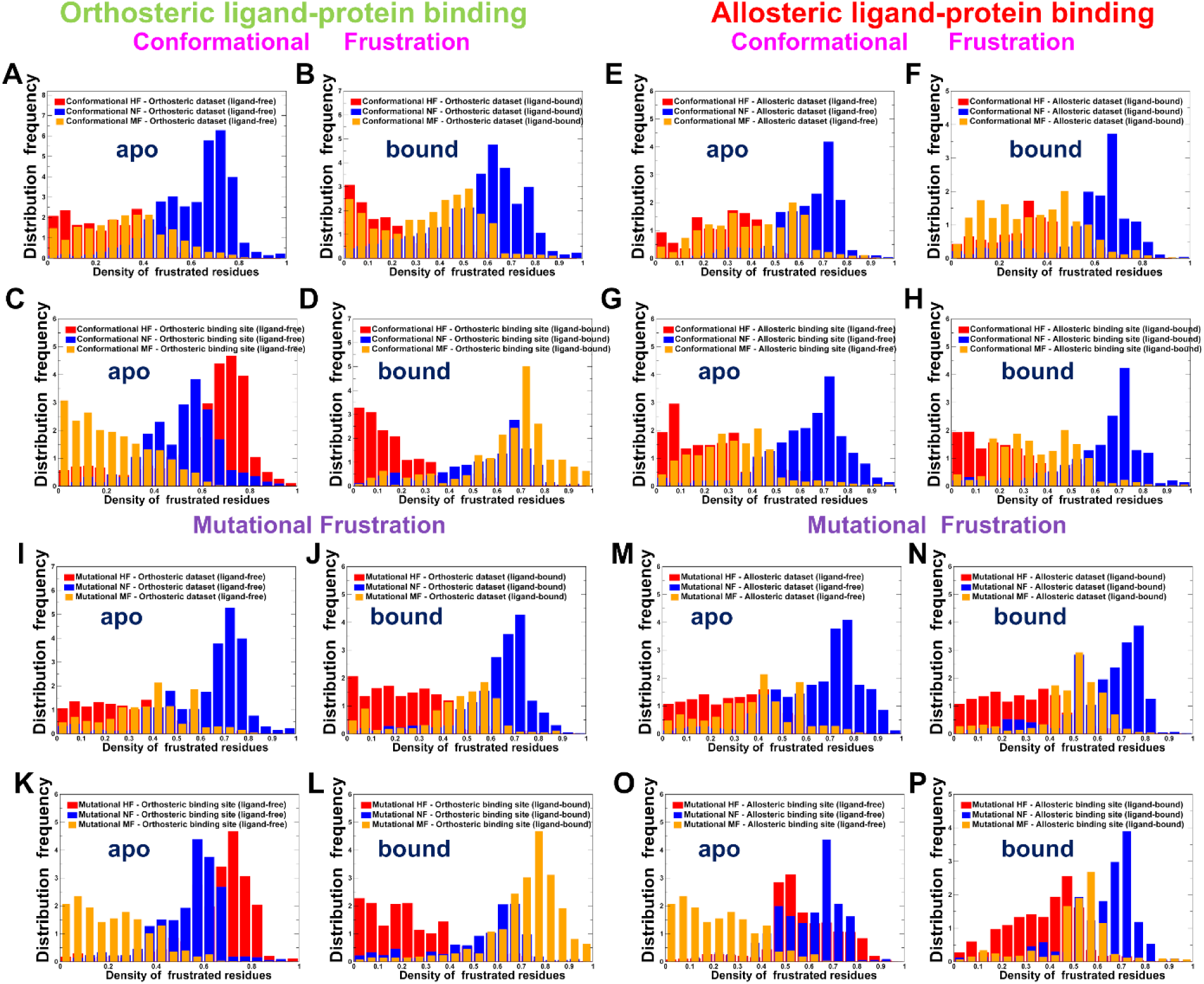
Frustration dichotomy encodes the energetic grammar of orthosteric versus allosteric binding sites. (A–D) Conformational frustration profiles for orthosteric complexes. (A) Global protein frustration in the apo state. (B) Global protein frustration in the holo state. (C) Binding-site frustration in the apo state. (D) Binding-site frustration in the holo state. Orthosteric sites undergo pronounced frustration quenching upon ligand binding: highly frustrated contacts (magenta, Z < −1.0) decrease from 28% in the apo state, while minimally frustrated contacts (orange, Z > 0.78) increase to 64% in the holo state. Global fold frustration remains largely unchanged. (E–H) Conformational frustration profiles for allosteric complexes. (E) Global protein frustration in the apo state. (F) Global protein frustration in the holo state. (G) Binding-site frustration in the apo state. (H) Binding-site frustration in the holo state. In contrast to orthosteric sites, neutral frustration (red, −1.0 ≤ Z ≤ 0.78) dominates both apo (71%) and holo (68%) allosteric sites with no systematic shift upon ligand binding. (I–L) Mutational frustration profiles for orthosteric complexes. (I) Global mutational frustration in the apo state. (J) Global mutational frustration in the holo state. (K) Orthosteric binding-site mutational frustration in the apo state. (L) Orthosteric binding-site mutational frustration in the holo state. Orthosteric sites show strong enrichment of minimal mutational frustration (79%, orange) upon ligand binding. (M–P) Mutational frustration profiles for allosteric complexes. (M) Global mutational frustration in the apo state. (N) Global mutational frustration in the holo state. (O) Allosteric binding-site mutational frustration in the apo state. (P) Allosteric binding-site mutational frustration in the holo state. Allosteric sites remain dominated by neutral mutational frustration (58%, red) in both states.

In all cases, minimally frustrated residues dominated the hydrophobic core (∼45–55%), highly frustrated residues localized to functional loops and interfaces (∼15–20%), and neutral frustration characterized surface-exposed and flexible regions (∼30–40%). These distributions were invariant between apo and holo states and did not differ between orthosteric and allosteric complexes, demonstrating that the predictive dichotomy does not originate from global fold organization. Instead, it must arise from the energetic architecture of the binding sites themselves and from how those sites respond to ligand binding. In the apo state, orthosteric binding sites are enriched in highly frustrated contacts as ∼ 28% of binding-site residues are highly frustrated compared to ∼12% in the global fold (Figure 4C). The remaining binding site residues are distributed between neutral (41%) and minimal (31%) frustration. Upon ligand binding, orthosteric sites undergo pronounced frustration quenching, generating the steep energetic funnels required for geometric convergence. In the holo state, the fraction of minimally frustrated residues increases from 31% to 64%, while highly frustrated contacts decrease from 28% to 8% (Figure 4D). Neutral frustration correspondingly decreases from 41% to 28%. Importantly, this reorganization is localized to the binding site, while global fold frustration remains essentially unchanged (Figure 4A–B).

Thus, ligand binding actively reshapes the local energy landscape rather than simply occupying a pre-existing cavity. More than one quarter of binding-site residues transition from strained or ambiguous energetic states into optimally stabilized interactions. This frustration quenching generates a strong directional gradient in conformational space—a deep energetic funnel that guides sampling toward a unique native geometry. Critically, this directional signal is created by ligand binding itself and is absent in the apo structure. AI models trained predominantly on holo complexes and optimized to detect stabilization patterns are therefore ideally positioned to exploit these ligand-induced funnels for precise pose placement at orthosteric sites.

Mutational frustration profiles reinforce the prevalence of neutral frustration for apo and holo forms of the global protein folds (Figure 4I,J) while revealing ligand-induced frustration quenching in orthosteric binding regions (Figure 4K,L). Orthosteric binding sites are strongly enriched in minimally frustrated residues, with 79% of binding-site positions classified as minimally frustrated in the holo state (Figure 4K,L). This pattern reflects tight purifying selection and dense coevolutionary coupling, generating the statistical signatures exploited by AI architectures for pocket identification. Orthosteric sites thus provide both energetic and evolutionary guidance for pocket localization and ligand pose placement.

### Persistent Neutrality and Evolutionary Constraints Prevent Energetic Convergence at Allosteric Sites and Reinforce Prediction Dichotomy

Allosteric binding sites in the apo state are dominated by neutral frustration, with 71% of residues classified as neutrally frustrated (Figure 4G). Highly frustrated contacts are rare, and minimally frustrated residues are sparse. This persistent neutrality means that allosteric binding regions lack distinctive energetic landmarks in the frustration landscape. In stark contrast to orthosteric complexes, allosteric binding sites exhibit no analogous frustration quenching upon ligand binding (Figure 4H). In both apo and holo states, neutral frustration dominates the binding region (71% and 68%, respectively), with minimally frustrated residues remaining modest (21% in apo, 24% in holo) and highly frustrated contacts rare (Figure 4G,H). The energetic frustration distribution of allosteric binding sites may be partially redistributed but displays persistent neutrality patterns upon ligand binding. As a result, multiple putative allosteric pockets can satisfy similar interaction topologies producing energetic degeneracy and preventing reproducible pocket detection and ligand pose convergence. Mutational frustration reveals a parallel divergence in evolutionary organization that compounds the prediction challenge of allosteric binding. Neutral mutational frustration is persistent not only for the global protein folds of allosteric complexes (Figure 4 M,N) but also dominates the frustration patterns of allosteric binding regions in both apo and holo states (Figure 4O,P) with only 34% of residues are minimally frustrated. This evolutionary permissiveness reflects the functional logic of allostery, which requires conformational and evolutionary plasticity to mediate large structural rearrangements through moderate energetic barriers. Allosteric ligand binding lacks strong coevolutionary signals for pocket detection and ligand-induced energetic funnels for pose convergence, producing a dual deficit in pocket detectability and geometric specificity.

### Structural Organization of Frustration Landscapes Encodes Binding-Site Predictability

The quantitative frustration patterns establish a clear energetic dichotomy between orthosteric and allosteric binding sites, but aggregate distributions cannot reveal how these differences manifest in three dimensions. Mapping frustration states onto representative protein structures provides this spatial context, revealing consistent architectural principles that govern pocket architecture and directly inform the dual prediction challenge of pocket identification and pose placement Orthosteric complexes exhibit a conserved spatial architecture that transcends individual protein families (Figure 5). Across the representative panel binding pockets are uniformly compact and deeply embedded within the protein core, typically at domain interfaces or within conserved catalytic clefts. Critically, minimally frustrated residues form extensive, spatially contiguous networks. In each complex, the minimally frustrated region envelops the binding pocket in a continuous shell that extends 8–12 Å into the surrounding structure, creating broad, cooperatively stabilized zones. This spatial continuity is the structural manifestation of the ligand-induced frustration quenching where 64% of binding-site residues transition to minimal frustration upon ligand engagement. The resulting landscape provides a distinctive energetic signal to AI models for unambiguous pocket identification, and an extended directional funnel that guides conformational sampling toward a unique native geometry. Residue-level distribution profiles of conformational frustration (Figure S5) and mutational frustration (Figure S6) for representative panel of orthosteric complexes corroborate this organization, revealing dense clusters of minimally frustrated residues at positions surrounding the binding pocket which is the recurrent biophysical signature that pattern-recognition architectures are optimized to detect. Notably, the spatial extent of these networks correlates with prediction confidence: systems with the most extensive minimal frustration shells correspond to those where AF3 and Chai-1 achieve sub-angstrom accuracy.

**Figure 5.**
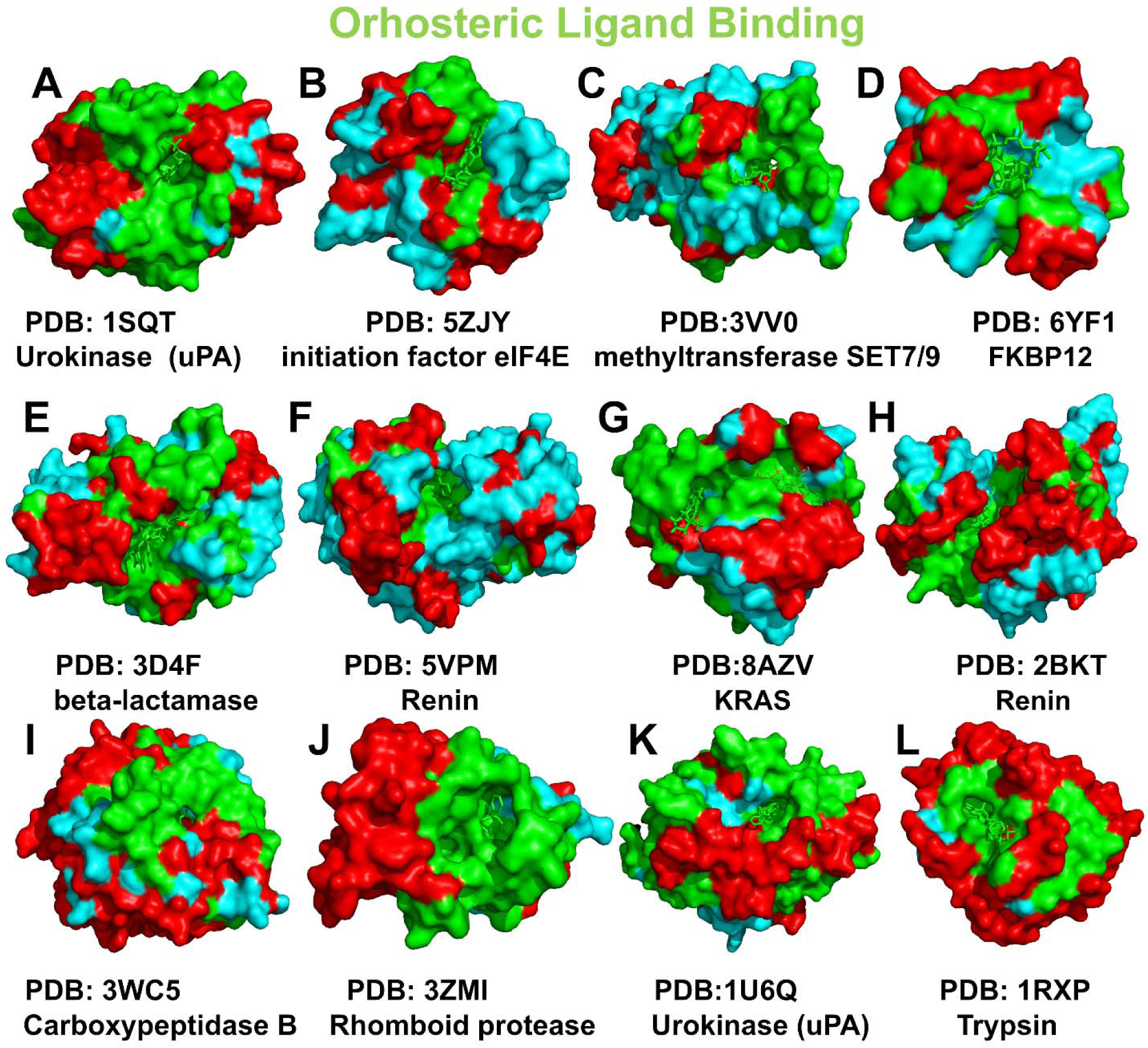
Structural maps of orthosteric complexes reveal extensive minimally frustrated networks across diverse protein families. For a representative panel of orthosteric ligand–protein complexes minimally frustrated regions are shown as green surfaces, neutrally frustrated regions as red surfaces. Ligands are shown as green sticks. Orthosteric pockets are uniformly small, deeply buried clefts. Minimal frustration extends well beyond the immediate binding pocket into neighboring structural elements, creating broad, connected stabilization zones whose spatial extent correlates with prediction confidence.

Allosteric complexes display a fundamentally distinct spatial grammar that holds across diverse regulatory architectures (Figure 6). The representative panel includes buried allosteric pockets, surface-exposed allosteric grooves, and cryptic sites that form upon ligand binding. Despite this architectural diversity, the frustration signature is remarkably consistent. Neutral frustration dominates throughout the protein fold, including directly within binding regions. Minimal frustration appears only as isolated, spatially disconnected patches that are scattered within expansive neutrally frustrated zones. In deeply buried allosteric pockets that superficially resemble orthosteric sites, the frustration maps reveal the critical distinction: these pockets lack the extended minimal frustration networks characteristic of their orthosteric counterparts. The surrounding region remains predominantly neutral, with only sporadic minimal frustration at key anchor points. For surface-exposed sites, neutral frustration pervades the entire binding interface, with minimal frustration relegated to distal structural elements uninvolved in ligand recognition. Residue-level distribution profiles of conformational frustration (Figure S7) and mutational frustration (Figure S8) for representative panel of allosteric complexes validate this organization, showing sparse, non-contiguous minimal frustration clusters and pervasive neutral mutational frustration that confirms evolutionary permissiveness. The absence of spatially coherent minimal frustration networks has two direct consequences for prediction. First, binding regions lack distinctive energetic landmarks, complicating reliable pocket identification. Second, even when pockets are approximately localized, the absence of an extended energetic funnel prevents geometric convergence to a unique pose. The isolated minimal frustration patches may provide weak local constraints, anchoring specific interactions without specifying global orientation, but they are insufficient to resolve the full three-dimensional arrangement. Models encounter multiple energetically comparable conformations without a dominant gradient to resolve ambiguity.

**Figure 6.**
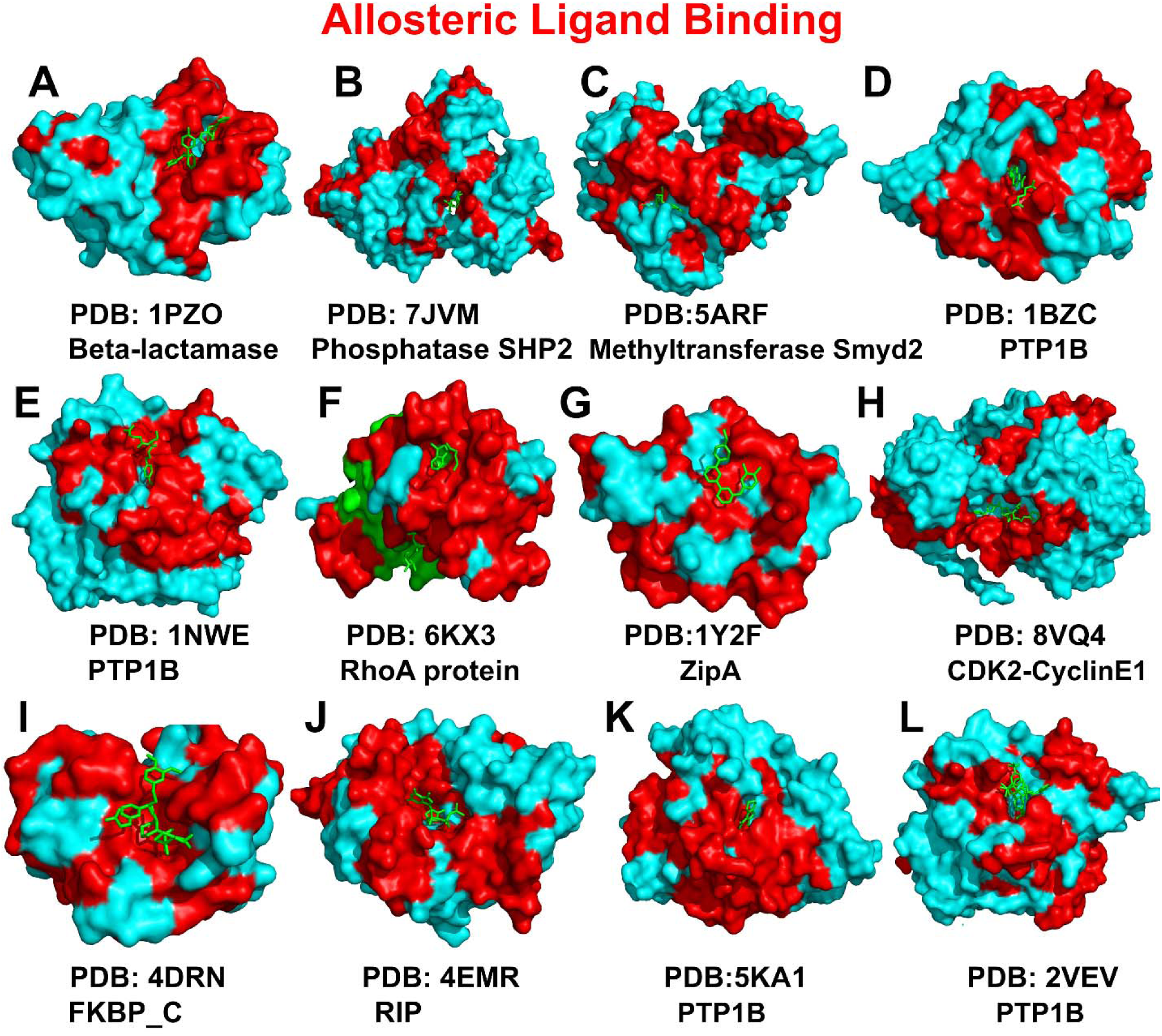
Structural maps of allosteric complexes show neutral frustration dominating binding site environments across diverse regulatory architectures. For a representative panel of allosteric complexes neutral frustration (red surfaces) dominates both global folds and binding site regions. Allosteric pockets—whether deeply buried or surface-exposed—are embedded within extensive neutrally frustrated zones, with only isolated, disconnected patches of minimal frustration (green) providing weak local constraints. Ligands (green sticks) sit within these neutrally frustrated environments, reflecting the absence of directional energetic gradients.

Together, these spatial features encode the dual challenge of allosteric prediction. Orthosteric landscapes broadcast a spatially coherent, evolutionarily reinforced signal through connected minimal frustration networks—a signature that specifies a unique solution by providing both location and orientation. Allosteric landscapes, by contrast, remain neutrally frustrated, energetically degenerate, and evolutionarily permissive in both apo and holo states, withholding the directional signals AI architectures require. Prediction accuracy thus becomes a quantitative reporter of energy landscape architecture, transforming model performance from a descriptive statistic into mechanistic insight into the physical grammar of allosteric recognition.

## Discussion

The integration of large-scale AI benchmarking with energy landscape analysis reveals a fundamental principle: the predictability of a binding site is not an architectural accident but is encoded in its biophysical organization. Across five state-of-the-art co-folding architectures—each embodying distinct design principles—the pattern is remarkably consistent. Orthosteric predictions achieve near-experimental accuracy with unimodal convergence, while allosteric predictions collapse systematically into broad, flat distributions. Mean errors double from 2–4 Å to 5–7 Å, success rates plummet from above 80% to below 35%. This uniformity across diverse inductive biases argues against explanations rooted in training data or architectural idiosyncrasies. It points instead to a common underlying constraint: the energetic landscapes of allosteric sites simply do not provide the signals current AI architectures are equipped to exploit. The nature of this limitation becomes clear in the decoupling between contact topology recovery and geometric precision. Allosteric predictions achieve moderately high QS-scores (0.70–0.85) even when RMSD exceeds 5 Å, indicating that models correctly identify which residues should interact—they recover the vocabulary of allosteric interactions. However, AI models cannot resolve the syntax of allosteric binding (the precise three-dimensional arrangement) because the landscape itself offers no unique solution. This is the signature of energetic degeneracy where multiple geometrically distinct conformations can satisfy similar interaction topologies in the absence of a steep local funnel. Local frustration analysis provides the physical basis for this dichotomy. In orthosteric sites, ligand binding actively reorganizes the local energy landscape, converting 28% of binding-site residues from highly frustrated in the apo state to minimally frustrated in the holo state. This frustration quenching creates steep directional gradients that funnel conformational sampling toward a unique geometry. The extensive, connected minimal frustration networks surrounding orthosteric pockets offer a spatial manifestation of these funnels. Allosteric sites, by contrast, maintain 68–71% neutral frustration in both apo and holo states, with no systematic reorganization upon ligand binding. Neutral frustration dominates, and minimal frustration appears only as isolated, disconnected patches. Mutational frustration reveals contrasting evolutionary constraints that compound the allosteric challenge. Orthosteric sites show overwhelming dominance of minimal mutational frustration (79%), indicating strong purifying selection that generates rich coevolutionary signals. Allosteric sites exhibit elevated neutral mutational frustration (58%), reflecting evolutionary permissiveness that erodes the sequence-based signals language models exploit. The results demonstrate that AI prediction accuracy serves as a quantitative reporter of underlying biophysical organization. Orthosteric sites broadcast strong signals through ligand-induced funnels, evolutionary constraints, and spatially coherent minimal frustration networks. Allosteric ligand binding is characterized by persistent neutral frustration, mutational permissiveness, and conformational heterogeneity. The allosteric blind spot, in this light, may reflect the glimpse of biophysical grammar of allostery that conceals a unique solution and challenges memorization patterns which AI models are trained to decipher. The systematic limitations of AI co-folding models on allosteric complexes, coupled with the energetic concealment revealed by frustration analysis, resonate with emerging recent work interrogating the limits of AI for protein-ligand prediction. Three complementary studies released very recently, each addressing distinct aspects of co-folding performance, provide striking conceptual alignment with our findings and provides compelling external validation for the biophysical framework we have established.^87–89^ In a prospective evaluation of 557 SARS-CoV-2 Mac1-ligand complexes Kim et al. demonstrated that while AF3, Boltz-2, and Chai-1 each reproduced >50% of poses within 2 Å RMSD, all models failed to recapitulate essential protein conformational changes required for ligand binding.^87^ This inability to model conformational adaptation directly parallels the systematic failure we observe on allosteric complexes, where structural plasticity is not incidental but definitional. The PLINDER resource introduced by Durairaj et al. addresses the foundational challenge of rigorous, leakage-minimized benchmarking, providing over 449,000 protein-ligand systems with multi-level similarity metrics that enable realistic assessment of model generalization.^88^ Their recognition that diverse inference scenarios, including novel pockets and protein-ligand combinations, require specialized stratified test sets directly supports our demonstration that allosteric sites represent a fundamental generalization challenge rather than a simple performance decrement. The diversity of allosteric site architectures suggests that different allosteric mechanisms may present different prediction challenges; larger datasets with finer mechanistic annotation could enable stratification of allosteric sites by their frustration signatures and corresponding prediction outcomes. Most directly, Zhang et al. benchmarked co-folding methods on 218 covalent complexes and found that while models achieved superior accuracy on familiar systems, performance declined markedly for novel pocket-ligand pairs^89^ which is a pattern mirroring the allosteric performance gap we report in this study.

In conclusion, our study establishes a principled, biophysics-informed explainable AI framework that transforms the allosteric blind spot from limitation into insight, providing both a diagnostic tool for evaluating next-generation models and a concrete roadmap for developing energy-aware architectures capable of navigating nature’s most sophisticated regulatory landscapes.

## Methods

### Orthosteric Datasets

The MDT test set served as the primary orthosteric reference, comprising 412 complexes^39^. Structural challenge was enforced by requiring AlphaFold-predicted apo-pocket deviation from experimental holo-conformation >2.0 Å RMSD or pocket LDDT <0.80. Ligands were restricted to drug-like molecules (200–650 Da), excluding fragments and macrocycles^39^, with no more than ten complexes from any single study. The datasets were filtered to include only complexes deposited in the Protein Data Bank after 2020, ensuring they post-date model training sets. Complexes with >30% sequence identity to training data were excluded (verified via PDB70 and MMseqs2 clustering). To expand statistical power, we incorporated the PLOC subset from a comprehensive structural dataset^86^ containing 1,863 structures with 2,325 binding pockets and 1,647 unique ligands binding to orthosteric sites across 76 Pfam families. Resolution ≤3.0 Å. Combined with MDT, the comprehensive orthosteric reference totaled 2,275 complexes.

### Allosteric Datasets

Kinase-specific allosteric complexes were derived from the Kinase Conformation Resource (KinCoRe)^83,84^ a geometry-driven classification system using DFG motif and αC-helix conformation. Type III inhibitors bind adjacent to the ATP site (distance >6.0 Å from hinge region, minimum three contacts with back-pocket residues). Type IV inhibitors bind >6.5 Å from both hinge region and conserved αC-helix Glu(+4) residue. From this resource, we extracted 217 Type III/IV complexes, augmented with a curated X-ray crystallography repository^85^ containing 136 allosteric kinase inhibitors (resolution ≤3.5 Å, MW ≥250 Da, no primary site occupancy). The kinase allosteric dataset totals 353 complexes. To capture allosteric diversity, we incorporated the PLA subset^86^, containing 1,613 structures with 1,830 binding pockets and 1,277 unique ligands spanning 83 Pfam families across >500 organisms. Resolution ≤3.2 Å. All complexes were experimentally validated as authentic allosteric binders with sites spatially distinct from orthosteric pockets. The combined allosteric ensemble totals 1,966 complexes.

### AI Co-folding Architectures

Five state-of-the-art architectures were benchmarked following protocols adapted from PoseBusters.^31^ All models received identical inputs: protein sequences from PDB structures, ligands in SMILES format. For each complex, 25 independent predictions were generated (10 recycling steps, 200 diffusion steps, 5 seeds, 5 diffusion samples per seed) on NVIDIA A100 40GB GPUs. The ligand-protein chain-pair iPTM score was used for ranking. Scripts are available in the accompanying code repository. AF3 employs a diffusion-based architecture operating on a unified “cloud of atoms,” simultaneously diffusing protein and ligand coordinates.^30^ Predictions were generated with templates enabled (default) and without templates (AF3-NT). Confidence metrics include pTM, ipTM, per-atom pLDDT, and PAE. Protenix is an open-source implementation of AF3 using unified tokenization of polymers and small molecules while maintaining architectural fidelity.^34^ Boltz-2 extends diffusion-based co-folding with affinity and binder-classification heads trained on experimental and MD-derived data.^35^ It incorporates prediction heads for affinity estimation and binder classification, using confidence-guided sampling to prioritize predictions with high internal consistency Chai-1 uses ESM-2 PLM embeddings with paired attention mechanisms, incorporating evolutionary information directly into structure prediction.^38^ Predictions were made with the ESM embedding option enabled. DynamicBind employs an SE(3)-equivariant diffusion graph network that iteratively adjusts receptor structure to accommodate the ligand.^39^ Training involves progressively distorting holo structures with noise and learning the reverse diffusion process that recovers native conformations. During inference, DynamicBind generates 20 candidate trajectories per target, scored using a composite metric combining LDDT scores and predicted binding affinities.

### Evaluation Metrics and Alignment Pipeline

All predicted protein structures were superimposed onto experimental references using TM-align^90^ for sequence-independent structural alignment. Several ligand RMSD tools such as LigRMSD^91^, DockRMSD^92^ and spyRMSD^93^ which are commonly applied in docking evaluations were initially employed but large scale pipeline execution of diverse AI models require clean ligand definitions, consistent atom ordering, and stable residue naming conventions which appear to be highly heterogeneous and inconsistent across different AI models outputs. OpenStructure framework was adopted in the final protocol which allows direct control over structure alignment, ligand extraction, and RMSD calculation within a single framework.^94^ We calculated the ligand pose RMSD with the OpenStructure toolkit (OST) using the Symmetry-Corrected RMSD Scorer (SCRMSDScorer) which enumerates symmetry-equivalent atom assignments and returns the lowest RMSD.^94^ The binding pocket was defined as all protein residues with at least one atom within 8 Å of any reference ligand atom. After defining the pocket, RMSD was calculated over the backbone atoms of these residues following the same global structural alignment used for ligand RMSD. QS-score quantified recovery of native contact topology independent of pose geometry, measuring the fraction of native ligand–protein contacts (4.5 Å cutoff) correctly recovered. LDDT-LP (Local Distance Difference Test for Ligand-Protein interfaces) evaluated preservation of local atomic distances within the binding pocket, with sensitivity to side-chain interactions. Ligand-centric confidence metrics included L-pLDDT (mean pLDDT over ligand atoms) and L-PAE (average PAE involving ligand atoms) for AF3/Chai-1, and ipTM for Boltz-2. Confidence-Error Correlation was assessed by mapping confidence scores against actual RMSD.

### Local Frustration Analysis

Local frustration analysis^73–78^ quantified energetic optimality of native interactions by comparing them to decoy ensembles. Two complementary indices were computed for every residue pair in both apo and holo states. Conformational frustration assesses sensitivity to local structural perturbations. For each contact, 1,000 decoy configurations are generated by randomizing residue identities and interatomic distances within native contact geometry. The native energy is compared to the decoy distribution; Z-scores below −1.0 indicate energetic strain or conformational conflict, while scores above 0.78 indicate strong optimization. Mutational frustration evaluates evolutionary optimality by comparing native interaction energy to 1,000 computational mutations at the same positions (all possible amino acid substitutions with fixed backbone geometry). Z-scores above 0.78 indicate native pairing is more favorable than most alternatives, reflecting evolutionary optimization. Both frustration indices were expressed as Z-scores, defined as:

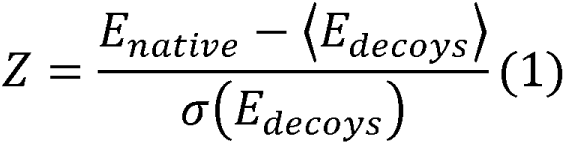

Following validated thresholds^73–75^, contacts were classified as minimally frustrated (Z > 0.78), highly frustrated (Z < −1.0), or neutrally frustrated (−1.0 ≤ Z ≤ 0.78). Residue-level profiles were generated by computing local density of frustrated contacts within a 5 Å radius. A residue was assigned a dominant frustration category if >50% of its interacting contacts belonged to the same class. Analysis was performed at two levels: across entire complexes for global landscape architecture, and within binding-site regions (residues within 4 Å of ligand).

### Quantification and Statistical Analysis

For each model, the best prediction per complex (lowest ligand pose RMSD) was selected from generated models. Average RMSD values represent the mean of best-run RMSDs across all systems in each dataset. Statistical significance of differences between orthosteric and allosteric predictions was assessed using paired two-tailed t-tests (p < 0.05) with Benjamini–Hochberg FDR correction.^95^ Residue-level frustration densities were compared using Mann–Whitney U test, with effect sizes quantified by Cliff’s delta.^96^ Spearman’s rank correlation was computed between per-residue frustration Z-scores and per-residue model confidence scores.

### Software Implementation

All computations were performed in Python 3.10 using SciPy (v1.10) for hypothesis testing, statsmodels (v0.14) for multiple comparison correction, and custom scripts for effect size and correlation analysis. Visualizations were generated with Matplotlib (v3.7) and Seaborn (v0.12). The full evaluation pipeline, including all scripts for input preparation, model execution, output postprocessing, and statistical analysis, is available in our open-source repository with detailed documentation for community adoption and extension.

## Resource availability Lead contact

Further information and requests for resources and materials should be directed to and will be fulfilled by the lead contact, Professor Gennady M. Verkhivker

## Materials Availability

No materials were synthesized in this study.

## Data and Code Availability

All unprocessed data and original code that the paper reports are freely available in an online repository that meets the criteria for digital longevity, implementation of FAIR standards, and community support.

## Data

- All data have been deposited at ZENODO (https://zenodo.org/records/18424406) and are publicly available as of the date of publication at DOI <u>10.5281/zenodo.18424405</u>.

## Code

- All original code has been deposited at (https://github.com/Verkhivker-Research-Group/AI-CoFolding-Allostery-Benchmark) and is publicly available at DOI <u>10.5281/zenodo.18424405</u> as of the date of publication.

## Additional information

- Any additional information required to reanalyze the data reported in this paper is available from the lead contact upon request.

Data for this article, including description of data types, all data, software and scripts are freely available at the Github repository https://github.com/Verkhivker-Research-Group/AI-CoFolding-Allostery-Benchmark/tree/main and the Zenodo archive https://zenodo.org/records/18424406 Github repository https://github.com/Verkhivker-Research-Group/AI-CoFolding-Allostery-Benchmark/tree/main contains information about Orthosteric Dataset and Allosteric Dataset. All scripts required to reproduce the predictions reported here are publicly available in GitHub repository. The github repository contains processed prediction outputs, evaluation results, figures, and scripts used in the study. The Zenodo archive https://zenodo.org/records/18424406 provides the datasets, processed prediction outputs, evaluation results, and scripts used in the study. Protein and ligand structures are not included in this archive. All structures are publicly available from the RCSB Protein Data Bank (http://www.rcsb.org) and can be automatically downloaded using the provided scripts. The rendering of protein structures was done with UCSF ChimeraX package (https://www.rbvi.ucsf.edu/chimerax/) and Pymol (https://pymol.org/2/). All structures are publicly available from the RCSB Protein Data Bank and can be automatically downloaded using the provided scripts. The data supporting this article have also been included in the Supplementary Information.

## Acknowledgments

This research was funded by the National Institutes of Health under Award 1R01AI181600-01, 5R01AI181600-02 and Subaward 6069-SC24-11 to G.V. G.V acknowledges support from Schmid College of Science and Technology at Chapman University for providing computing resources at the Keck Center for Science and Engineering.

## Author Contributions

Conceptualization, G.V.; Methodology, V.P., B.F., W.G., M.L., L.T., G.V.; Software, V.P., B.F., W.G., M.L., L.T., G.V.; Validation, V.P., B.F., W.G., M.L., L.T., G.V. ; Formal analysis, V.P., B.F., W.G., M.L., L.T., G.V.; Investigation, V.P., B.F., W.G., M.L., L.T., G.V.; Resources, V.P., B.F., W.G., M.L., L.T., G.V.; Data curation, V.P., B.F., W.G., M.L., L.T., G.V.; Writing—original draft preparation, V.P., G.V. ; Writing—review and editing, V.P., G.V. ; Visualization, V.P., B.F., W.G., M.L., L.T., G.V.; Supervision G.V.; Project administration, G.V. ; Funding acquisition, G.V.

All authors have read and agreed to their individual contributions. All authors have approved the submission of this manuscript, and it has been submitted exclusively to Cell Reports Physical Science.

## References

(1) Jumper, J.; Evans, R.; Pritzel, A.; Green, T.; Figurnov, M.; Ronneberger, O.; Tunyasuvunakool, K.; Bates, R.; Žídek, A.; Potapenko, A.; Bridgland, A.; Meyer, C.; Kohl, S. A. A.; Ballard, A. J.; Cowie, A.; Romera-Paredes, B.; Nikolov, S.; Jain, R.; Adler, J.; Back, T.; Petersen, S.; Reiman, D.; Clancy, E.; Zielinski, M.; Steinegger, M.; Pacholska, M.; Berghammer, T.; Bodenstein, S.; Silver, D.; Vinyals, O.; Senior, A. W.; Kavukcuoglu, K.; Kohli, P.; Hassabis, D. Highly Accurate Protein Structure Prediction with AlphaFold. Nature 2021, 596, 583–589. doi: 10.1038/s41586-021-03819-2.

(2) Tunyasuvunakool, K.; Adler, J.; Wu, Z.; Green, T.; Zielinski, M.; Žídek, A.; Bridgland, A.; Cowie, A.; Meyer, C.; Laydon, A.; Velankar, S.; Kleywegt, G. J.; Bateman, A.; Evans, R.; Pritzel, A.; Figurnov, M.; Ronneberger, O.; Bates, R.; Kohl, S. A. A.; Potapenko, A.; Ballard, A. J.; Romera-Paredes, B.; Nikolov, S.; Jain, R.; Clancy, E.; Reiman, D.; Petersen, S.; Senior, A. W.; Kavukcuoglu, K.; Birney, E.; Kohli, P.; Jumper, J.; Hassabis, D. Highly Accurate Protein Structure Prediction for the Human Proteome. Nature 2021, 596, 590–596. doi: 10.1038/s41586-021-03828-1.

(3) Chen, H.; Engkvist, O.; Wang, Y.; Olivecrona, M.; Blaschke, T. The rise of deep learning in drug discovery. Drug Discov Today 2018, 23, 1241–1250. doi: 10.1016/j.drudis.2018.01.039.

(3) Merchant, A.; Batzner, S.; Schoenholz, S. S.; Aykol, M.; Cheon, G.; Cubuk, E. D. Scaling Deep Learning for Materials Discovery. Nature 2023, 624, 80–85. doi: 10.1038/s41586-023-06735-9.

(4) Madani, A.; Krause, B.; Greene, E. R.; Subramanian, S.; Mohr, B. P.; Holton, J. M.; Olmos, J. L., Jr.; Xiong, C.; Sun, Z. Z.; Socher, R.; Fraser, J. S.; Naik, N. Large Language Models Generate Functional Protein Sequences across Diverse Families. Nat Biotechnol. 2023 , 41,1099–1106. doi: 10.1038/s41587-022-01618-2.

(5) Zheng, Y.; Koh, H. Y.; Ju, J.; Yang, M.; May, L. T.; Webb, G. I.; Li, L.; Pan, S.; Church, G. Large Language Models for Drug Discovery, and Development. Patterns. 2025, 6, 101346. doi: 10.1016/j.patter.2025.101346.

(6) Evans, R.; O’Neill, M.; Pritzel, A.; Antropova, N.; Senior, A.; Green, T.; Žídek, A.; Bates, R.; Blackwell, S.; Yim, J.; Ronneberger, O.; Bodenstein, S.; Zielinski, M.; Bridgland, A.; Potapenko, A.; Cowie, A.; Tunyasuvunakool, K.; Jain, R.; Clancy, E.; Kohli, P.; Jumper, J.; Hassabis, D. Protein Complex Prediction with AlphaFold-Multimer. bioRxiv 2021. doi: 10.1101/2021.10.04.463034

(7) Lin, Z.; Akin, H.; Rao, R.; Hie, B.; Zhu, Z.; Lu, W.; Smetanin, N.; Verkuil, R.; Kabeli, O.; Shmueli, Y.; dos Santos Costa, A.; Fazel-Zarandi, M.; Sercu, T.; Candido, S.; Rives, A. Evolutionary-Scale Prediction of Atomic-Level Protein Structure with a Language Model. Science 2023, 379, 1123–1130. doi: 10.1126/science.ade2574.

(8) Wu, R.; Ding, F.; Wang, R.; Shen, R.; Zhang, X.; Luo, S.; Su, C.; Wu, Z.; Xie, Q.; Berger, B.; Ma, J.; Peng, J. High-Resolution de Novo Structure Prediction from Primary Sequence, bioArxiv, 2022. 10.1101/2022.07.21.500999.

(9) Baek, M.; DiMaio, F.; Anishchenko, I.; Dauparas, J.; Ovchinnikov, S.; Lee, G. R.; Wang, J.; Cong, Q.; Kinch, L. N.; Schaeffer, R. D.; Millán, C.; Park, H.; Adams, C.; Glassman, C. R.; DeGiovanni, A.; Pereira, J. H.; Rodrigues, A. V.; van Dijk, A. A.; Ebrecht, A. C.; Opperman, D. J.; Sagmeister, T.; Buhlheller, C.; Pavkov-Keller, T.; Rathinaswamy, M. K.; Dalwadi, U.; Yip, C. K.; Burke, J. E.; Garcia, K. C.; Grishin, N. V.; Adams, P. D.; Read, R. J.; Baker, D. Accurate Prediction of Protein Structures and Interactions Using a Three-Track Neural Network. Science 2021, 373, 871–876. 10.1126/science.abj8754.

(10) Krishna, R.; Wang, J.; Ahern, W.; Sturmfels, P.; Venkatesh, P.; Kalvet, I.; Lee, G. R.; Morey-Burrows, F. S.; Anishchenko, I.; Humphreys, I. R.; McHugh, R.; Vafeados, D.; Li, X.; Sutherland, G. A.; Hitchcock, A.; Hunter, C. N.; Kang, A.; Brackenbrough, E.; Bera, A. K.; Baek, M.; DiMaio, F.; Baker, D. Generalized Biomolecular Modeling and Design with RoseTTAFold All-Atom. Science 2024, 384 (6693), eadl2528. doi: 10.1126/science.adl2528.

(11) Watson, J. L.; Juergens, D.; Bennett, N. R.; Trippe, B. L.; Yim, J.; Eisenach, H. E.; Ahern, W.; Borst, A. J.; Ragotte, R. J.; Milles, L. F.; Wicky, B. I. M.; Hanikel, N.; Pellock, S. J.; Courbet, A.; Sheffler, W.; Wang, J.; Venkatesh, P.; Sappington, I.; Torres, S. V.; Lauko, A.; De Bortoli, V.; Mathieu, E.; Ovchinnikov, S.; Barzilay, R.; Jaakkola, T. S.; DiMaio, F.; Baek, M.; Baker, D. De Novo Design of Protein Structure and Function with RFdiffusion. Nature 2023, 620, 1089–1100. 10.1038/s41586-023-06415-8.

(12) Fleishman, S.J., Horovitz, A. Extending the New Generation of Structure Predictors to Account for Dynamics and Allostery. J. Mol. Biol. 2021, 433, 167007. doi: 10.1016/j.jmb.2021.167007.

(13) Del Alamo, D.; Sala, D.; Mchaourab, H.S.; Meiler, J. Sampling Alternative Conformational States of Transporters and Receptors with AlphaFold2. Elife 2022, 11, e75751. doi: 10.7554/eLife.75751.

(14) Stein, R. A.; Mchaourab, H. S. SPEACH_AF: Sampling Protein Ensembles and Conformational Heterogeneity with Alphafold2. PLoS Comput Biol. 2022, 18, e1010483. doi: 10.1371/journal.pcbi.1010483.

(15) Saldaño, T.; Escobedo, N.; Marchetti, J.; Zea, D. J.; Mac Donagh, J.; Velez Rueda, A. J.; Gonik, E.; García Melani, A.; Novomisky Nechcoff, J.; Salas, M. N.; Peters, T.; Demitroff, N.; Fernandez Alberti, S.; Palopoli, N.; Fornasari, M. S.; Parisi, G. Impact of Protein Conformational Diversity on AlphaFold Predictions. Bioinformatics 2022, 38, 2742–2748. doi: 10.1093/bioinformatics/btac202.

(16) Chakravarty, D.; Porter, L. L. AlphaFold2 Fails to Predict Protein Fold Switching. Protein Sci. 2022 , 31, e4353. doi: 10.1002/pro.4353.

(17) Chakravarty, D.; Schafer, J. W.; Chen, E. A.; Thole, J. F.; Ronish, L. A.; Lee, M.; Porter, L. L. AlphaFold Predictions of Fold-Switched Conformations Are Driven by Structure Memorization. Nat Commun. 2024 ,15, 7296. doi: 10.1038/s41467-024-51801-z.

(18) Wayment-Steele, H. K.; Ovchinnikov, S.; Colwell, L.; Kern, D. Predicting multiple conformations via sequence clustering and AlphaFold2. Nature 2023. doi: 10.1038/s41586-023-06832-9.

(19) Porter, L. L.; Chakravarty, D.; Schafer, J. W.; Chen, E. A. ColabFold Predicts Alternative Protein Structures from Single Sequences, Coevolution Unnecessary for AF-Cluster. bioRxiv, 2023. doi: 10.1101/2023.11.21.567977.

(20) Bryant, P.; Noé, F. Structure Prediction of Alternative Protein Conformations. Nat Commun. 2024 , 15, 7328. doi: 10.1038/s41467-024-51507-2.

(21) Ahdritz, G.; Bouatta, N.; Floristean, C.; Kadyan, S.; Xia, Q.; Gerecke, W.; O’Donnell, T. J.; Berenberg, D.; Fisk, I.; Zanichelli, N.; Zhang, B.; Nowaczynski, A.; Wang, B.; Stepniewska-Dziubinska, M. M.; Zhang, S.; Ojewole, A.; Guney, M. E.; Biderman, S.; Watkins, A. M.; Ra, S.; Lorenzo, P. R.; Nivon, L.; Weitzner, B.; Ban, Y.-E. A.; Chen, S.; Zhang, M.; Li, C.; Song, S. L.; He, Y.; Sorger, P. K.; Mostaque, E.; Zhang, Z.; Bonneau, R.; AlQuraishi, M. OpenFold: Retraining AlphaFold2 Yields New Insights into Its Learning Mechanisms and Capacity for Generalization. Nat Methods. 2024, 21, 1514–1524. doi: 10.1038/s41592-024-02272-z.

(22) Li, Z.; Liu, X.; Chen, W.; Shen, F.; Bi, H.; Ke, G.; Zhang, L. Uni-Fold: An Open-Source Platform for Developing Protein Folding Models beyond AlphaFold, bioRxiv 2022. 10.1101/2022.08.04.502811.

(23) Raisinghani, N.; Alshahrani, M.; Gupta, G.; Tian, H.; Xiao, S.; Tao, P.; Verkhivker, G. M. Integration of a Randomized Sequence Scanning Approach in AlphaFold2 and Local Frustration Profiling of Conformational States Enable Interpretable Atomistic Characterization of Conformational Ensembles and Detection of Hidden Allosteric States in the ABL1 Protein Kinase. J. Chem. Theory Comput. 2024, 20, 5317–5336. doi: 10.1021/acs.jctc.4c00222.

(24) Raisinghani, N.; Alshahrani, M.; Gupta, G.; Verkhivker, G. Predicting Mutation-Induced Allosteric Changes in Structures and Conformational Ensembles of the ABL Kinase Using AlphaFold2 Adaptations with Alanine Sequence Scanning. Int J Mol Sci. 2024 , 25, 10082. doi: 10.3390/ijms251810082.

(25) Guan, X.; Tang, Q.-Y.; Ren, W.; Chen, M.; Wang, W.; Wolynes, P. G.; Li, W. Predicting Protein Conformational Motions Using Energetic Frustration Analysis and AlphaFold2. Proc. Natl. Acad. Sci. U.S.A. 2024, 121, e2410662121. doi: 10.1073/pnas.2410662121

(26) Xu, M.; Yu, L.; Song, Y.; Shi, C.; Ermon, S.; Tang, J. GeoDiff: A Geometric Diffusion Model for Molecular Conformation Generation. arXiv 2022. Doi: 10.48550/ARXIV.2203.02923.

(27) Jing, B.; Corso, G.; Chang, J.; Barzilay, R.; Jaakkola, T. Torsional Diffusion for Molecular Conformer Generation. arXiv 2022. Doi: 10.48550/ARXIV.2206.01729.

(28) Corso, G.; Stärk, H.; Jing, B.; Barzilay, R.; Jaakkola, T. DiffDock: Diffusion Steps, Twists, and Turns for Molecular Docking. arXiv 2022. Doi; 10.48550/ARXIV.2210.01776.

(29) Corso, G.; Deng, A.; Fry, B.; Polizzi, N.; Barzilay, R.; Jaakkola, T. Deep Confident Steps to New Pockets: Strategies for Docking Generalization. arXiv 2024. Doi: 10.48550/ARXIV.2402.18396.

(30) Abramson, J.; Adler, J.; Dunger, J.; Evans, R.; Green, T.; Pritzel, A.; Ronneberger, O.; Willmore, L.; Ballard, A. J.; Bambrick, J.; Bodenstein, S. W.; Evans, D. A.; Hung, C.-C.; O’Neill, M.; Reiman, D.; Tunyasuvunakool, K.; Wu, Z.; Žemgulytė, A.; Arvaniti, E.; Beattie, C.; Bertolli, O.; Bridgland, A.; Cherepanov, A.; Congreve, M.; Cowen-Rivers, A. I.; Cowie, A.; Figurnov, M.; Fuchs, F. B.; Gladman, H.; Jain, R.; Khan, Y. A.; Low, C. M. R.; Perlin, K.; Potapenko, A.; Savy, P.; Singh, S.; Stecula, A.; Thillaisundaram, A.; Tong, C.; Yakneen, S.; Zhong, E. D.; Zielinski, M.; Žídek, A.; Bapst, V.; Kohli, P.; Jaderberg, M.; Hassabis, D.; Jumper, J. M. Accurate Structure Prediction of Biomolecular Interactions with AlphaFold 3. Nature 2024 , 630, 493–500. doi: 10.1038/s41586-024-07487-w.

(31) Buttenschoen, M.; Morris, G. M.; Deane, C. M. PoseBusters: AI-Based Docking Methods Fail to Generate Physically Valid Poses or Generalise to Novel Sequences. Chem. Sci. 2024, 15, 3130–3139. 10.1039/d3sc04185a.

(32) Trott, O.; Olson, A. J. AutoDock Vina: Improving the Speed and Accuracy of Docking with a New Scoring Function, Efficient Optimization, and Multithreading. J Comput Chem. 2010 31, 455–61. doi: 10.1002/jcc.21334.

(33) Masters, M. R.; Mahmoud, A. H.; Lill, M. A. Investigating Whether Deep Learning Models for Co-Folding Learn the Physics of Protein-Ligand Interactions. Nat Commun. 2025 , 16, 8854. doi: 10.1038/s41467-025-63947-5.

(34) Chen, X.; Zhang, Y.; Lu, C.; Ma, W.; Guan, J.; Gong, C.; Yang, J.; Zhang, H.; Zhang, K.; Wu, S.; Zhou, K.; Yang, Y.; Liu, Z.; Wang, L.; Shi, B.; Shi, S.; Xiao, W. Protenix – Advancing Structure Prediction Through a Comprehensive AlphaFold3 Reproduction. bioRxiv 2025. doi: 10.1101/2025.01.08.631967.

(35) Passaro, S.; Corso, G.; Wohlwend, J.; Reveiz, M.; Thaler, S.; Somnath, V. R.; Getz, N.; Portnoi, T.; Roy, J.; Stark, H.; Kwabi-Addo, D.; Beaini, D.; Jaakkola, T.; Barzilay, R. Boltz-2: Towards Accurate and Efficient Binding Affinity Prediction. bioRxiv 2025 doi: 10.1101/2025.06.14.659707.

(36) Tosstorff, A.; Rudolph, M. G.; Benz, J.; Kuhn, B.; Kramer, C.; Sharpe, M.; Huang, C.; Metz, A.; Hazemann, J.; Ritz, D.; Sweeney, A. M.; Gilson, M. K. The CASP 16 Experimental Protein-Ligand Datasets. Proteins 2026 ,94, 79–85. doi: 10.1002/prot.70053.

(37) Gilson, M. K.; Eberhardt, J.; Škrinjar, P.; Durairaj, J.; Robin, X.; Kryshtafovych, A. Assessment of Pharmaceutical Protein–Ligand Pose and Affinity Predictions in CASP16. Proteins 2026, 94, 249–266. doi: 10.1002/prot.70061.

(38) Boitreaud, J.; Dent, J.; McPartlon, M.; Meier, J.; Reis, V.; Rogozhnikov, A.; Wu, K. Chai-1: Decoding the Molecular Interactions of Life. bioRxiv 2024. doi: 10.1101/2024.10.10.615955.

(39) Lu, W.; Zhang, J.; Huang, W.; Zhang, Z.; Jia, X.; Wang, Z.; Shi, L.; Li, C.; Wolynes, P. G.; Zheng, S. DynamicBind: Predicting Ligand-Specific Protein-Ligand Complex Structure with a Deep Equivariant Generative Model. Nat Commun. 2024, 15, 1071. doi: 10.1038/s41467-024-45461-2.

(40) Stärk, H.; Ganea, O.-E.; Pattanaik, L.; Barzilay, R.; Jaakkola, T. EquiBind: Geometric Deep Learning for Drug Binding Structure Prediction. arXiv 2022. doi: 10.48550/ARXIV.2202.05146.

(41) Schneuing, A.; Harris, C.; Du, Y.; Didi, K.; Jamasb, A.; Igashov, I.; Du, W.; Gomes, C.; Blundell, T. L.; Lio, P.; Welling, M.; Bronstein, M.; Correia, B. Structure-Based Drug Design with Equivariant Diffusion Models. Nat Comput Sci. 2024, 4, 899–909. doi: 10.1038/s43588-024-00737-x.

(42) Lu, W.; Wu, Q.; Zhang, J.; Rao, J.; Li, C.; Zheng, S. TANKBind: Trigonometry-Aware Neural NetworKs for Drug-Protein Binding Structure Prediction. bioRxiv 2022. doi: 10.1101/2022.06.06.495043

(43) Dauparas, J.; Anishchenko, I.; Bennett, N.; Bai, H.; Ragotte, R. J.; Milles, L. F.; Wicky, B. I. M.; Courbet, A.; de Haas, R. J.; Bethel, N.; Leung, P. J. Y.; Huddy, T. F.; Pellock, S.; Tischer, D.; Chan, F.; Koepnick, B.; Nguyen, H.; Kang, A.; Sankaran, B.; Bera, A. K.; King, N. P.; Baker, D. Robust Deep Learning–Based Protein Sequence Design Using ProteinMPNN. Science 2022, 378, 49–56. doi: 10.1126/science.add2187.

(44) Hayes, T.; Rao, R.; Akin, H.; Sofroniew, N. J.; Oktay, D.; Lin, Z.; Verkuil, R.; Tran, V. Q.; Deaton, J.; Wiggert, M.; Badkundri, R.; Shafkat, I.; Gong, J.; Derry, A.; Molina, R. S.; Thomas, N.; Khan, Y. A.; Mishra, C.; Kim, C.; Bartie, L. J.; Nemeth, M.; Hsu, P. D.; Sercu, T.; Candido, S.; Rives, A. Simulating 500 Million Years of Evolution with a Language Model. Science 2025, 387, 850–858. doi: 10.1126/science.ads0018.

(45) Ingraham, J. B.; Baranov, M.; Costello, Z.; Barber, K. W.; Wang, W.; Ismail, A.; Frappier, V.; Lord, D. M.; Ng-Thow-Hing, C.; Van Vlack, E. R.; Tie, S.; Xue, V.; Cowles, S. C.; Leung, A.; Rodrigues, J. V.; Morales-Perez, C. L.; Ayoub, A. M.; Green, R.; Puentes, K.; Oplinger, F.; Panwar, N. V.; Obermeyer, F.; Root, A. R.; Beam, A. L.; Poelwijk, F. J.; Grigoryan, G. Illuminating Protein Space with a Programmable Generative Model. Nature 2023, 623, 1070–1078. doi: 10.1038/s41586-023-06728-8.

(46) Le Guilloux, V.; Schmidtke, P.; Tuffery, P. Fpocket: An Open Source Platform for Ligand Pocket Detection. BMC Bioinform. 2009, 10, 168. doi: 10.1186/1471-2105-10-168.

(47) Ghersi, D.; Sanchez, R. EasyMIFS and SiteHound: a toolkit for the identification of ligand-binding sites in protein structures. Bioinformatics 2009, 25, 3185–3186. doi: 10.1093/bioinformatics/btp562.

(48) Liu, Y.; Grimm, M.; Dai, W.; Hou, M.; Xiao, Z.-X.; Cao, Y. CB-Dock: A Web Server for Cavity Detection-Guided Protein–Ligand Blind Docking. Acta Pharmaco.l Sin. 2020, 41,138–144. doi: 10.1038/s41401-019-0228-6.

(49) Zhang, Z.; Li, Y.; Lin, B.; Schroeder, M.; Huang, B. Identification of Cavities on Protein Surface Using Multiple Computational Approaches for Drug Binding Site Prediction. Bioinformatics 2011, 27, 2083–2088. doi: 10.1093/bioinformatics/btr331.

(50) Krivák, R.; Hoksza, D. P2Rank: Machine Learning Based Tool for Rapid and Accurate Prediction of Ligand Binding Sites from Protein Structure. J. Cheminform. 2018, 10, 39. doi: 10.1186/s13321-018-0285-8.

(51) Polák, L.; Škoda, P.; Riedlová, K.; Krivák, R.; Novotný, M.; Hoksza, D. PrankWeb 4: A Modular Web Server for Protein–Ligand Binding Site Prediction and Downstream Analysis. Nucleic Acids Res. 2025, 53, W466–W471. doi: 10.1093/nar/gkaf421.

(52) Jian, J.-W.; Elumalai, P.; Pitti, T.; Wu, C. Y.; Tsai, K.-C.; Chang, J.-Y.; Peng, H.-P.; Yang, A.-S. Predicting Ligand Binding Sites on Protein Surfaces by 3-Dimensional Probability Density Distributions of Interacting Atoms. PLoS One 2016, 11, e0160315. doi: 10.1371/journal.pone.0160315.

(53) Jiménez, J.; Doerr, S.; Martínez-Rosell, G.; Rose, A. S.; De Fabritiis, G. DeepSite: Protein-Binding Site Predictor Using 3D-Convolutional Neural Networks. Bioinformatics 2017, 33, 3036–3042. 10.1093/bioinformatics/btx350.

(54) Jiang, M.; Wei, Z.; Zhang, S.; Wang, S.; Wang, X.; Li, Z. FRSite: Protein Drug Binding Site Prediction Based on Faster R–CNN. J Mol Graph Model. 2019, 93, 107454. doi: 10.1016/j.jmgm.2019.107454.

(55) Stepniewska-Dziubinska, M. M.; Zielenkiewicz, P.; Siedlecki, P. Improving Detection of Protein-Ligand Binding Sites with 3D Segmentation. Sci Rep. 2020, 10, 5035. doi: 10.1038/s41598-020-61860-z.

(56) Kozlovskii, I. Popov, P. Spatiotemporal identification of druggable binding sites using deep learning. Commun Biol. 2020 , 3, 618. doi: 10.1038/s42003-020-01350-0.

(57) Škrhák, V.; Novotný, M.; Feidakis, C. P.; Krivák, R.; Hoksza, D. CryptoBench: Cryptic Protein–Ligand Binding Sites Dataset and Benchmark. Bioinformatics 2024, 41, btae745. doi: 10.1093/bioinformatics/btae745.

(58) Sim, J.; Kim, D.; Kim, B.; Choi, J.; Lee, J. Recent Advances in AI-Driven Protein-Ligand Interaction Predictions. Curr.Opin Struct Biol. 2025, 92, 103020. doi: 10.1016/j.sbi.2025.103020.

(59) Rohulia, A.; Wang, Y.; Cheng, J. A Survey of Deep Learning Methods and Tools for Protein Binding Site Prediction. Methods Mol Biol. 2025, 2947, 89–108. doi: 10.1007/978-1-0716-4662-5_5.

(60) Carbery, A.; Buttenschoen, M.; Skyner, R.; von Delft, F.; Deane, C. M. Learnt Representations of Proteins Can Be Used for Accurate Prediction of Small Molecule Binding Sites on Experimentally Determined and Predicted Protein Structures. J Cheminform. 2024 16, 32. doi: 10.1186/s13321-024-00821-4.

(61) Smith, Z.; Strobel, M.; Vani, B. P.; Tiwary, P. Graph Attention Site Prediction (GrASP): Identifying Druggable Binding Sites Using Graph Neural Networks with Attention. J. Chem. Inf. Model. 2024, 64, 2637–2644. doi: 10.1021/acs.jcim.3c01698.

(62) Hoksza, D.; Gamouh, H. Exploration of Protein Sequence Embeddings for Protein-Ligand Binding Site Detection. 2022 IEEE International Conference on Bioinformatics and Biomedicine (BIBM), Las Vegas, NV, USA, 2022, pp. 3356–3361, doi: 10.1109/BIBM55620.2022.9995025.

(63) Gamouh, H.; Novotný, M.; Hoksza, D. Hybrid Protein–Ligand Binding Residue Prediction with Protein Language Models: Does the Structure Matter? Bioinformatics 2025 , 41, btaf431. doi: 10.1093/bioinformatics/btaf431.

(64) Schmirler, R.; Heinzinger, M.; Rost, B. Fine-Tuning Protein Language Models Boosts Predictions across Diverse Tasks. Nat Commun. 2024, 15, 7407. doi: 10.1038/s41467-024-51844-2.

(65) Greener, J. G.; Sternberg, M. J. AlloPred: Prediction of Allosteric Pockets on Proteins Using Normal Mode Perturbation Analysis. BMC Bioinform. 2015, 16, 335. doi: 10.1186/s12859-015-0771-1.

(66) Huang, W.; Lu, S.; Huang, Z.; Liu, X.; Mou, L.; Luo, Y.; Zhao, Y.; Liu, Y.; Chen, Z.; Hou, T.; Zhang, J. Allosite: A Method for Predicting Allosteric Sites. Bioinformatics, 2013, 29,2357–2359. doi: 10.1093/bioinformatics/btt399.

(67) Song, K.; Liu, X.; Huang, W.; Lu, S.; Shen, Q.; Zhang, L.; Zhang, J. Improved Method for the Identification and Validation of Allosteric Sites. J. Chem. Inf. Model. 2017, 57, 2358–2363. doi: 10.1021/acs.jcim.7b00014.

(68) He, J.; Liu, X.; Zhu, C.; Zha, J.; Li, Q.; Zhao, M.; Wei, J.; Li, M.; Wu, C.; Wang, J.; Jiao, Y.; Ning, S.; Zhou, J.; Hong, Y.; Liu, Y.; He, H.; Zhang, M.; Chen, F.; Li, Y.; He, X.; Wu, J.; Lu, S.; Song, K.; Lu, X.; Zhang, J. ASD2023: Towards the Integrating Landscapes of Allosteric Knowledgebase. Nucleic Acids Res. 2024, 52, D376–D383. doi: 10.1093/nar/gkad915.

(69) Khokhar, M.; Keskin, O.; Gursoy, A. DeepAllo: Allosteric Site Prediction Using Protein Language Modelwith Multitask Learning. Bioinformatics 2025, 41, btaf294. doi: 10.1093/bioinformatics/btaf294.

(70) Meller, A.; Ward, M.; Borowsky, J.; Kshirsagar, M.; Lotthammer, J. M.; Oviedo, F.; Ferres, J. L.; Bowman, G. R. Predicting Locations of Cryptic Pockets from Single Protein Structures Using the PocketMiner Graph Neural Network. Nat Commun. 2023, 14, 1177. doi: 10.1038/s41467-023-36699-3.

(71) Li, Y.; Yi, J.; Li, H.; Li, K.; Kang, F.; Deng, Y.; Wu, C.; Fu, X.; Jiang, D.; Cao, D. Decoding the Limits of Deep Learning in Molecular Docking for Drug Discovery. Chem. Sci. 2025, 16, 17374–17390. doi: 10.1039/d5sc05395a.

(72) La Sala, G.; Pfleger, C.; Käck, H.; Wissler, L.; Nevin, P.; Böhm, K.; Janet, J. P.; Schimpl, M.; Stubbs, C. J.; De Vivo, M.; Tyrchan, C.; Hogner, A.; Gohlke, H.; Frolov, A. I. Combining Structural and Coevolution Information to Unveil Allosteric Sites. Chem. Sci. 2023, 14, 7057–7067. doi: 10.1039/d2sc06272k.

(73) Ferreiro, D.U.; Hegler, J.A.; Komives, E.A.; Wolynes, P.G. Localizing frustration in native proteins and protein assemblies. Proc. Natl. Acad. Sci. U. S. A. 2007, 104, 19819–19824.

(74) Parra, R. G.; Schafer, N. P.; Radusky, L. G.; Tsai, M. Y.; Guzovsky, A. B.; Wolynes, P. G.; Ferreiro, D. U. Protein Frustratometer 2: A Tool to Localize Energetic Frustration in Protein Molecules, Now With Electrostatics. Nucleic Acids Res. 2016, 44 (W1), W356–360.

(75) Freiberger, M. I.; Wolynes, P. G.; Ferreiro, D. U.; Fuxreiter, M. Frustration in Fuzzy Protein Complexes Leads to Interaction Versatility. J Phys Chem B 2021, 125 (10), 2513–2520.

(76) Chen, M.; Chen, X.; Schafer, N.P.; Clementi, C.; Komives, E.A.; Ferreiro, D.U.; Wolynes, P.G. Surveying biomolecular frustration at atomic resolution. Nat. Commun. 2020, 11, 5944.

(77) Gianni, S.; Freiberger, M. I.; Jemth, P.; Ferreiro, D. U.; Wolynes, P. G.; Fuxreiter, M. Fuzziness and Frustration in the Energy Landscape of Protein Folding, Function, and Assembly. Acc. Chem. Res. 2021, 54 (5), 1251–1259.

(78) Parra, R. G.; Komives, E. A.; Wolynes, P. G.; Ferreiro, D. U. Frustration in Physiology and Molecular Medicine. Mol Aspects Med. 2025 103,101362. doi: 10.1016/j.mam.2025.101362.

(79) Dixit, A.; Verkhivker, G. M. The Energy Landscape Analysis of Cancer Mutations in Protein Kinases. PLoS ONE 2011, 6, e26071. doi: 10.1371/journal.pone.0026071.

(80) Verkhivker, G. M.; Agajanian, S.; Kassab, R.; Krishnan, K. Landscape-Based Protein Stability Analysis and Network Modeling of Multiple Conformational States of the SARS-CoV-2 Spike D614G Mutant: Conformational Plasticity and Frustration-Induced Allostery as Energetic Drivers of Highly Transmissible Spike Variants. J. Chem. Inf. Model. 2022, 62, 1956–1978. doi: 10.1021/acs.jcim.2c00124.

(81) Verkhivker, G. M.; Agajanian, S.; Kassab, R.; Krishnan, K. Frustration-Driven Allosteric Regulation and Signal Transmission in the SARS-CoV-2 Spike Omicron Trimer Structures: A Crosstalk of the Omicron Mutation Sites Allosterically Regulates Tradeoffs of Protein Stability and Conformational Adaptability. Phys. Chem. Chem. Phys. 2022, 24, 17723–17743. doi: 10.1039/d2cp01893d.

(82) Alshahrani, M.; Parikh, V.; Foley, B.; Verkhivker, G. Neutral Frustration Landscape Architecture of SARS-CoV-2 Spike-Antibody Interfaces Shapes Immune Evasion Mechanisms for Ultrapotent Neutralizing Antibodies and Determines Pathways of Viral Adaptation: Insights from Integrative Computational Approach. Phys. Chem. Chem. Phys 2026 DOI: 10.1039/D5CP04209G

(83) Modi, V.; Dunbrack, R. L., Jr. Defining a New Nomenclature for the Structures of Active and Inactive Kinases. Proc. Natl. Acad. Sci. U.S.A. 2019, 116, 6818–6827. doi: 10.1073/pnas.1814279116.

(84) Modi, V.; Dunbrack, R. L., Jr. Kincore: A Web Resource for Structural Classification of Protein Kinases and Their Inhibitors. Nucleic Acids Res. 2022, 50, D654–D664. doi: 10.1093/nar/gkab920.

(85) Hu, H.; Laufkötter, O.; Miljković, F.; Bajorath, J. Data Set of Competitive and Allosteric Protein Kinase Inhibitors Confirmed by X-Ray Crystallography. Data Brief. 2021, 35, 106816. doi: 10.1016/j.dib.2021.106816.

(86) Moine-Franel, A.; Mareuil, F.; Nilges, M.; Ciambur, C. B.; Sperandio, O. A Comprehensive Dataset of Protein-Protein Interactions and Ligand Binding Pockets for Advancing Drug Discovery. Sci Data. 2024 , 11, 402. doi: 10.1038/s41597-024-03233-z.

(87) Kim, J.; Correy, G. J.; Hall, B. W.; Rachman, M. M.; Mailhot, O.; Togo, T.; Gonciarz, R. L.; Jaishankar, P.; Neitz, R. J.; Hantz, E. R.; Doruk, Y. U.; Stevens, M. G. V.; Diolaiti, M. E.; Reid, R.; Gopalkrishnan, S.; Krogan, N. J.; Renslo, A. R.; Ashworth, A.; Shoichet, B. K.; Fraser, J. S. Large Scale Prospective Evaluation of Co-Folding across 557 Mac1-Ligand Complexes and Three Virtual Screens, bioRxiv. 2025. doi: 10.64898/2025.12.25.696505.

(88) Durairaj, J.; Adeshina, Y.; Cao, Z.; Zhang, X.; Oleinikovas, V.; Duignan, T.; McClure, Z.; Robin, X.; Studer, G.; Kovtun, D.; Rossi, E.; Zhou, G.; Veccham, S.; Isert, C.; Peng, Y.; Sundareson, P.; Akdel, M.; Corso, G.; Stärk, H.; Tauriello, G.; Carpenter, Z.; Bronstein, M.; Kucukbenli, E.; Schwede, T.; Naef, L. PLINDER: The Protein-Ligand Interactions Dataset and Evaluation Resource, 2024. 10.1101/2024.07.17.603955.

(89) Zhang, T.; Zhu, J.; Huang, Z.; Xie, J.; Pei, J.; Lai, L. Benchmarking Co-Folding Methods to Predict the Structures of Covalent Protein–Ligand Complexes. Acta Pharmacol. Sin. 2026. doi: 10.1038/s41401-025-01721-5.

(90) Zhang, Y. TM-Align: A Protein Structure Alignment Algorithm Based on the TM-Score. Nucleic Acids Res. 2005, 33, 2302–2309. doi: 10.1093/nar/gki524.

(91) Velázquez-Libera, J. L.; Durán-Verdugo, F.; Valdés-Jiménez, A.; Núñez-Vivanco, G.; Caballero, J. LigRMSD: A Web Server for Automatic Structure Matching and RMSD Calculations among Identical and Similar Compounds in Protein-Ligand Docking. Bioinformatics 2020, 36, 2912–2914. doi: 10.1093/bioinformatics/btaa018.

(92) Bell, E. W.; Zhang, Y. DockRMSD: An Open-Source Tool for Atom Mapping and RMSD Calculation of Symmetric Molecules through Graph Isomorphism. J Cheminform. 2019, 11, 40. doi: 10.1186/s13321-019-0362-7.

(93) Meli, R.; Biggin, P. C. Spyrmsd: Symmetry-Corrected RMSD Calculations in Python. J Cheminform. 2020, 12, 49. doi: 10.1186/s13321-020-00455-2.

(94) Biasini, M.; Schmidt, T.; Bienert, S.; Mariani, V.; Studer, G.; Haas, J.; Johner, N.; Schenk, A. D.; Philippsen, A.; Schwede, T. OpenStructure: An Integrated Software Framework for Computational Structural Biology. Acta Crystallogr. D Biol. Crystallogr. 2013, 69 , 701–709. doi: 10.1107/S0907444913007051.

(95) Benjamini, Y.; Hochberg, Y. Controlling the False Discovery Rate: A Practical and Powerful Approach to Multiple Testing. J. R. Stat. Soc. Ser. B Stat. Methodol. 1995, 57, 289–300. 10.1111/j.2517-6161.1995.tb02031.x.

(96) Chicco, D.; Sichenze, A.; Jurman, G. A Simple Guide to the Use of Student’s t-Test, Mann-Whitney U Test, Chi-Squared Test, and Kruskal-Wallis Test in Biostatistics. BioData Mining 2025, 18, 56. doi: 10.1186/s13040-025-00465-6.

