## Supplemental Files: Figures S1-S8 for "Decoding the Allosteric Paradox: A Dual Framework Integrating AI Cofolding Models with Landscape-Guided Interpretable AI Framework of Ligand-Protein Binding"

**Supplemental Information**
**Document S1. Figures S1–S8**

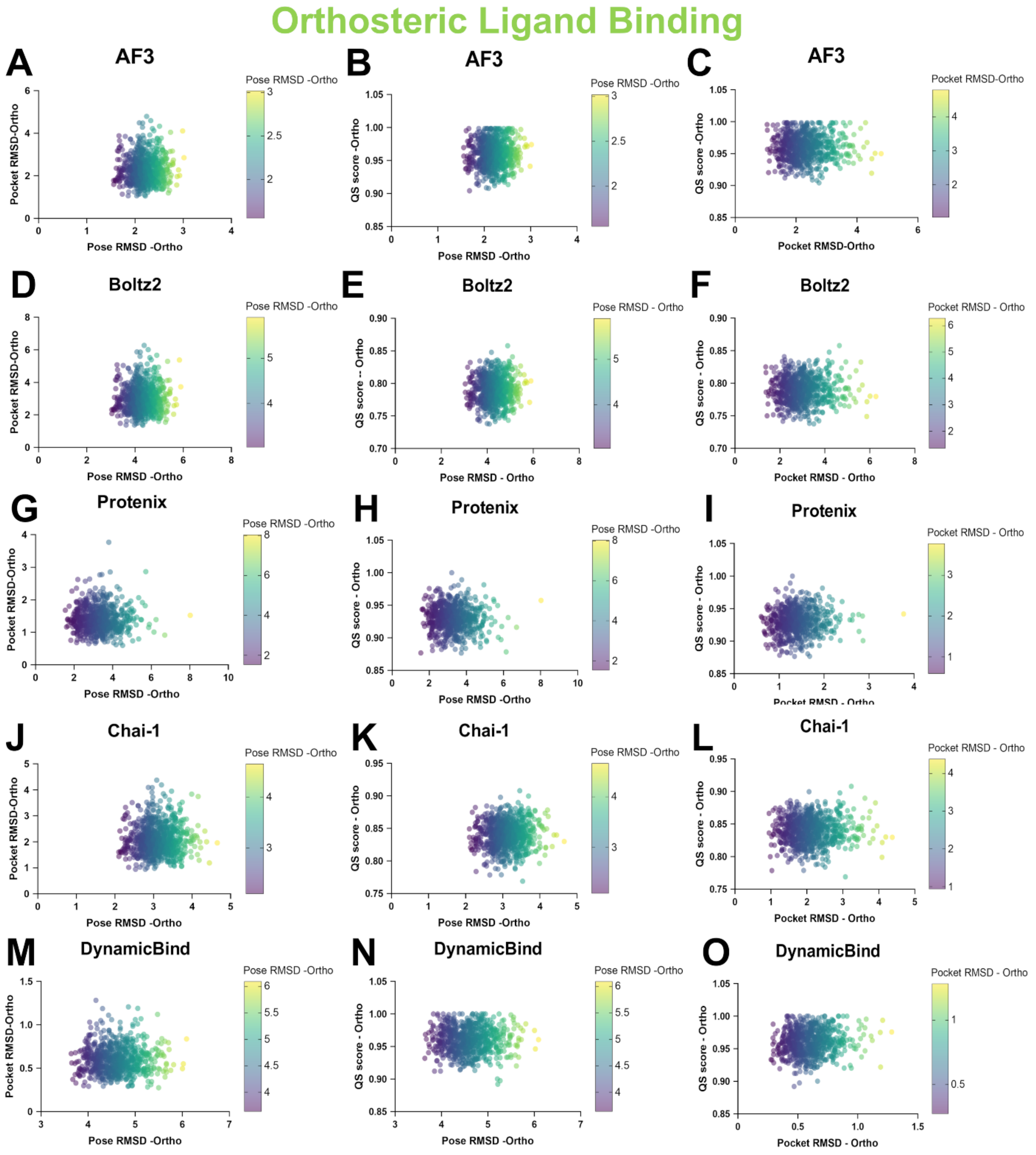


**Figure S1. Bulb scatter plots illustrating coupled geometric and topological accuracy for orthosteric complexes.** Bulb scatter plots show ligand pose RMSD versus pocket RMSD (left panels), ligand pose RMSD versus QS-score (middle panels) and pocket RMSD versus QS-score (right panels) for orthosteric complexes across all five AI models. Each point represents a predicted complex, with bulb size proportional to point density. In pose RMSD versus QS-score space, predictions cluster in the low-RMSD (<3 Å), high-QS (>0.90) quadrant, indicating strong coupling between geometric accuracy and native contact recovery. In pose RMSD versus pocket RMSD space, points concentrate near the origin, confirming that accurate pocket localization is tightly linked to correct pose placement. The compact, symmetric distributions reflect a well-defined energetic funnel that enforces convergence at orthosteric sites.

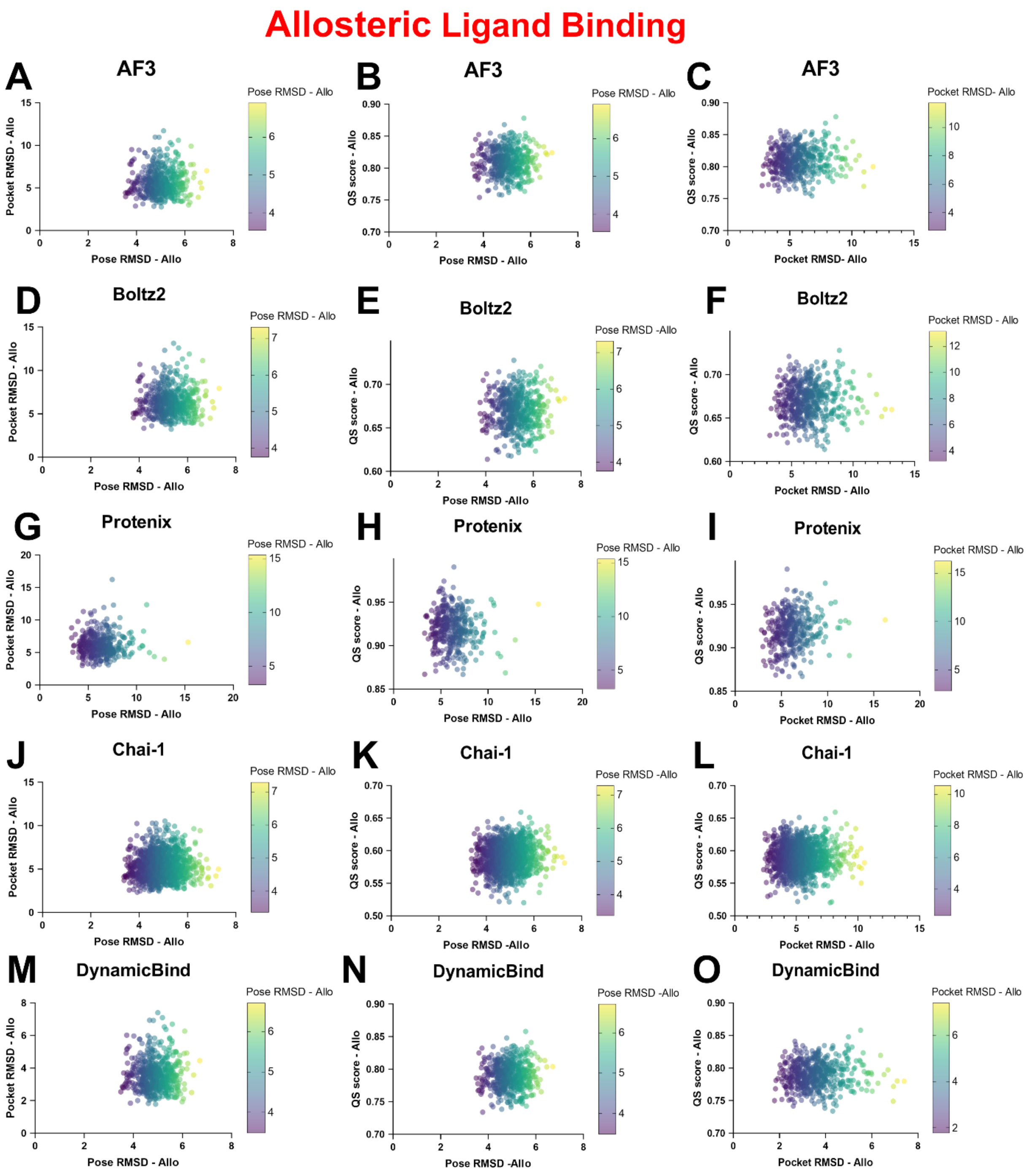


**Figure S2. Bulb scatter plots revealing decoupling between geometry and contact topology for allosteric complexes.** Bulb scatter plots show ligand pose RMSD versus pocket RMSD (left panels), ligand pose RMSD versus QS-score (middle panels) and pocket RMSD versus QS-score (right panels) for allosteric complexes across all five AI models. In pose RMSD versus pocket RMSD space, predictions show diagonal dispersion, with substantial density at low pocket RMSD (<3 Å) but high pose RMSD (>5 Å), indicating that pocket localization does not ensure correct pose placement. In pose RMSD versus QS-score space, QS-scores remain moderately high (≈0.6–0.9) over a wide RMSD range (<2 Å to >20 Å), demonstrating decoupling between interaction recovery and geometric precision. Together, these plots provide a visual signature of energetic degeneracy at allosteric sites, where multiple geometrically distinct solutions satisfy similar interaction topologies and prevent reproducible convergence.

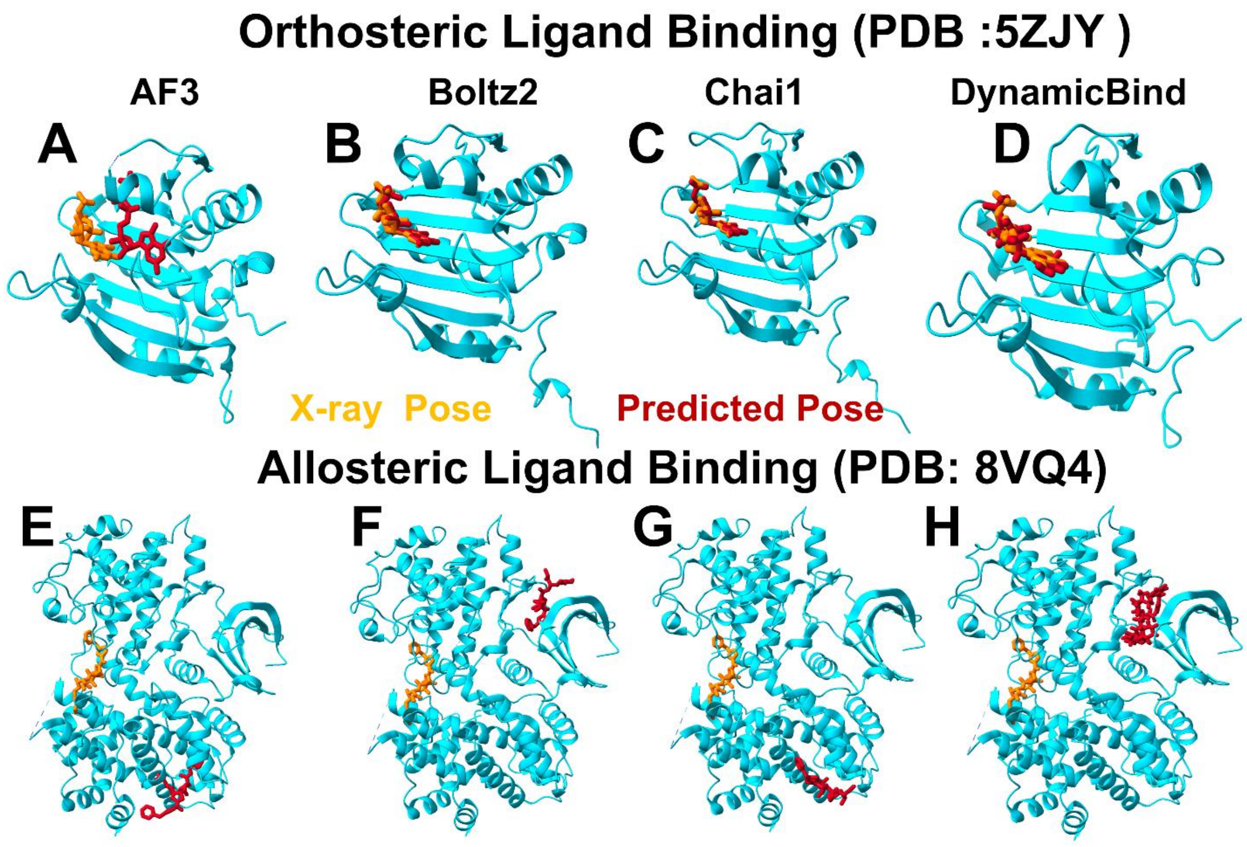


**Figure S3. Representative orthosteric and allosteric ligand predictions illustrate the performance dichotomy across AI architectures.** For each model—AF3, Boltz-2, Chai-1, and DynamicBind—two representative predictions are shown: one orthosteric complex (top row for each model) and one allosteric complex (bottom row for each model). The orthosteric complex is represented by the human eukaryotic initiation factor 4E (eIF4E) bound with the cap analogue ligand 5-methyl-GTP (MGP) at the classical cap binding site (pdb id 5ZJY). The allosteric complex is represented by a complex of cyclin dependent kinase 2 (CDK2) with its activator cyclin E1 in association with a small molecule inhibitor I-125A at a cryptic allosteric pocket at the CDK2-cyclin E1 interface (pdb id 8VQ4). In all panels, the experimental X-ray ligand pose is shown in green sticks, and the predicted ligand pose is shown in red sticks. Protein structures are shown in cartoon representation with binding-site residues highlighted. Orthosteric predictions closely reproduce the experimental geometry across all models, with near-perfect overlap between predicted and reference ligands with a notable exception of AF3 prediction. In contrast, allosteric predictions exhibit large geometric deviations: predicted ligands adopt alternative orientations, shift within the pocket, or fail to localize the correct binding site altogether. These examples visually illustrate the systematic divergence between high-fidelity orthosteric prediction and degraded allosteric performance observed quantitatively in Figures 2 and 3.

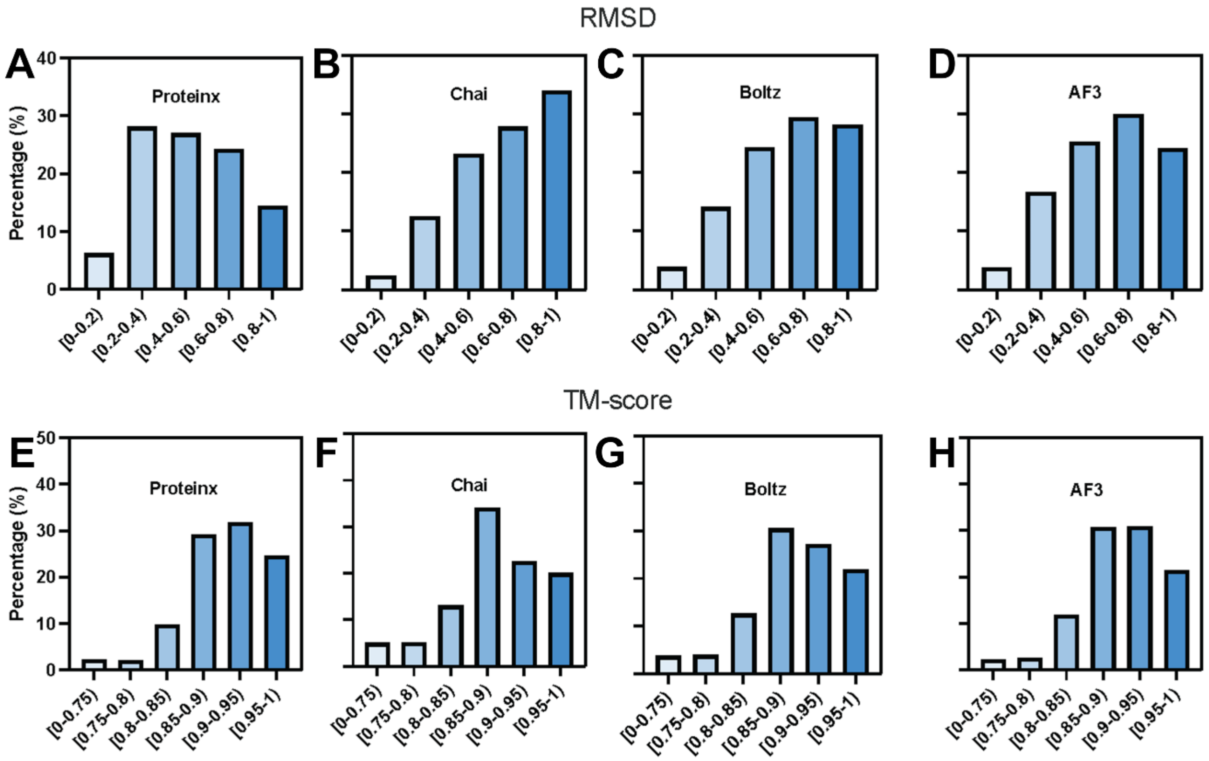


**Figure S4. Distribution of ultra-high-accuracy orthosteric predictions (ligand pose RMSD < 1.0 Å).** Histogram showing the number of orthosteric complexes achieving sub-angstrom ligand pose accuracy for each AI model. In the histograms, the bars follow a blue-toned monochromatic gradient that corresponds to the value bins on the x-axis (RMSDs for panels A-D and TM-score for panels E-H). The color mapping for each individual histogram is as follows: Lightest Blue is used for the bins representing the lowest values for RMSD and TM-score; Medium Blue for the middle-range bins; Darkest Blue for the bins representing the highest values for RMSD and TM-score. This hierarchy reflects architectural tradeoffs between maximal precision and robustness and highlights the capacity of current AI models to approach crystallographic accuracy when the energetic signal is strong, as in orthosteric binding sites.

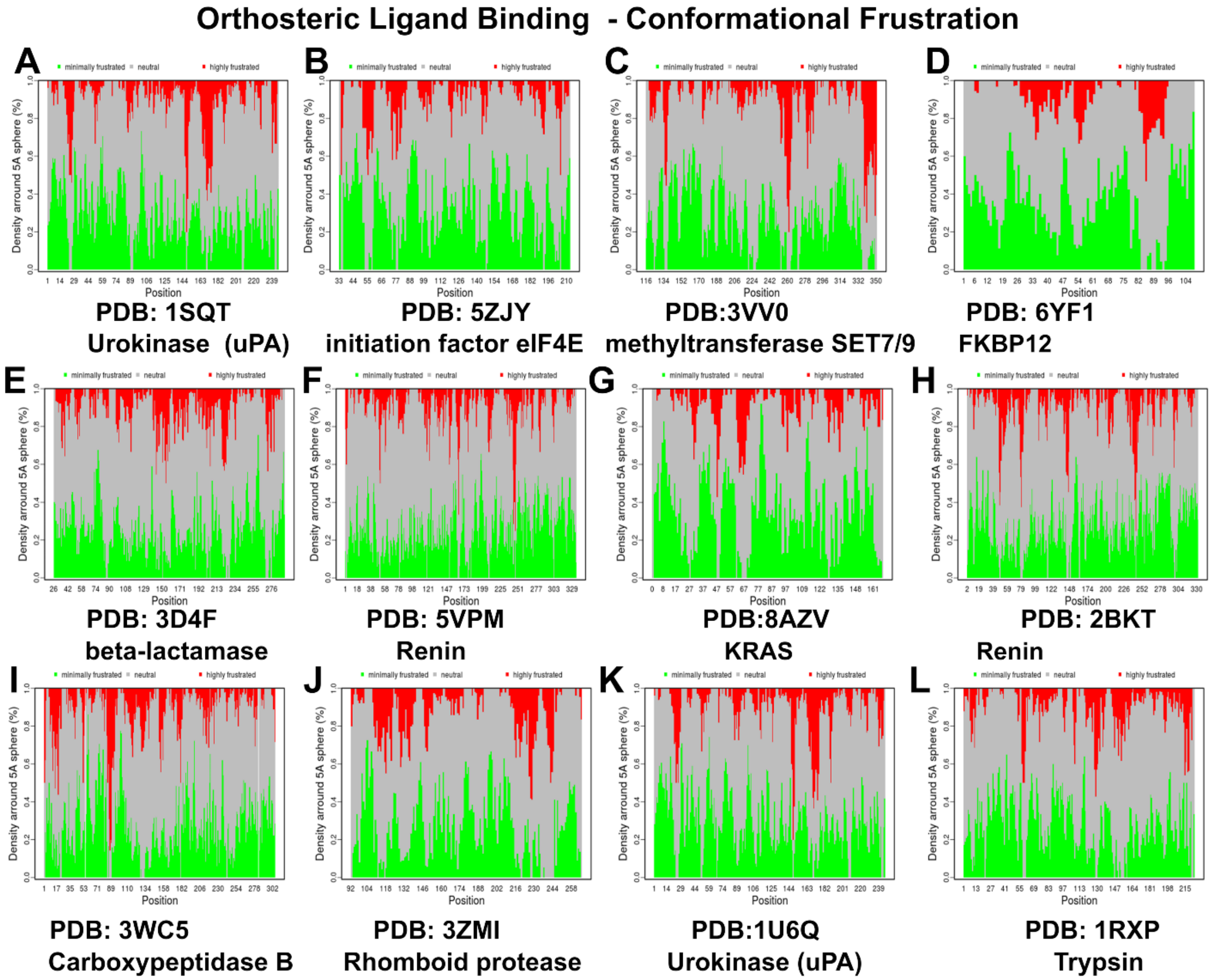


**Figure S5. Residue-level conformational frustration profiles for representative orthosteric ligand–protein complexes.** For each complex, the fraction of mutational frustration contact types is plotted per residue position: minimally frustrated (green), neutral (gray), and highly frustrated (red). Profiles are shown for (A) urokinase (uPA, PDB 1SQT), (B) initiation factor eIF4E (PDB 5ZJY), (C) methyltransferase SET7/9 (PDB 3VVO), (D) FKBP12 (PDB 6YF1), (E) beta-lactamase (PDB 3D4F), (F) renin (PDB 5VPM), (G) KRAS (PDB 8AZV), (H) renin (PDB 2BKT), (I) carboxypeptidase B (PDB 3WCI), (J) rhomboid protease (PDB 3ZMI), (K) urokinase (PDB 1U6Q), and (L) trypsin (PDB 1RXP). Across all orthosteric complexes, binding pockets exhibit pronounced enrichment of minimally frustrated residues forming spatially contiguous clusters that surround the ligand-binding site, reflecting strong local energetic stabilization and evolutionary optimization. Highly frustrated residues are located predominantly to flexible loop regions distal to the pocket.

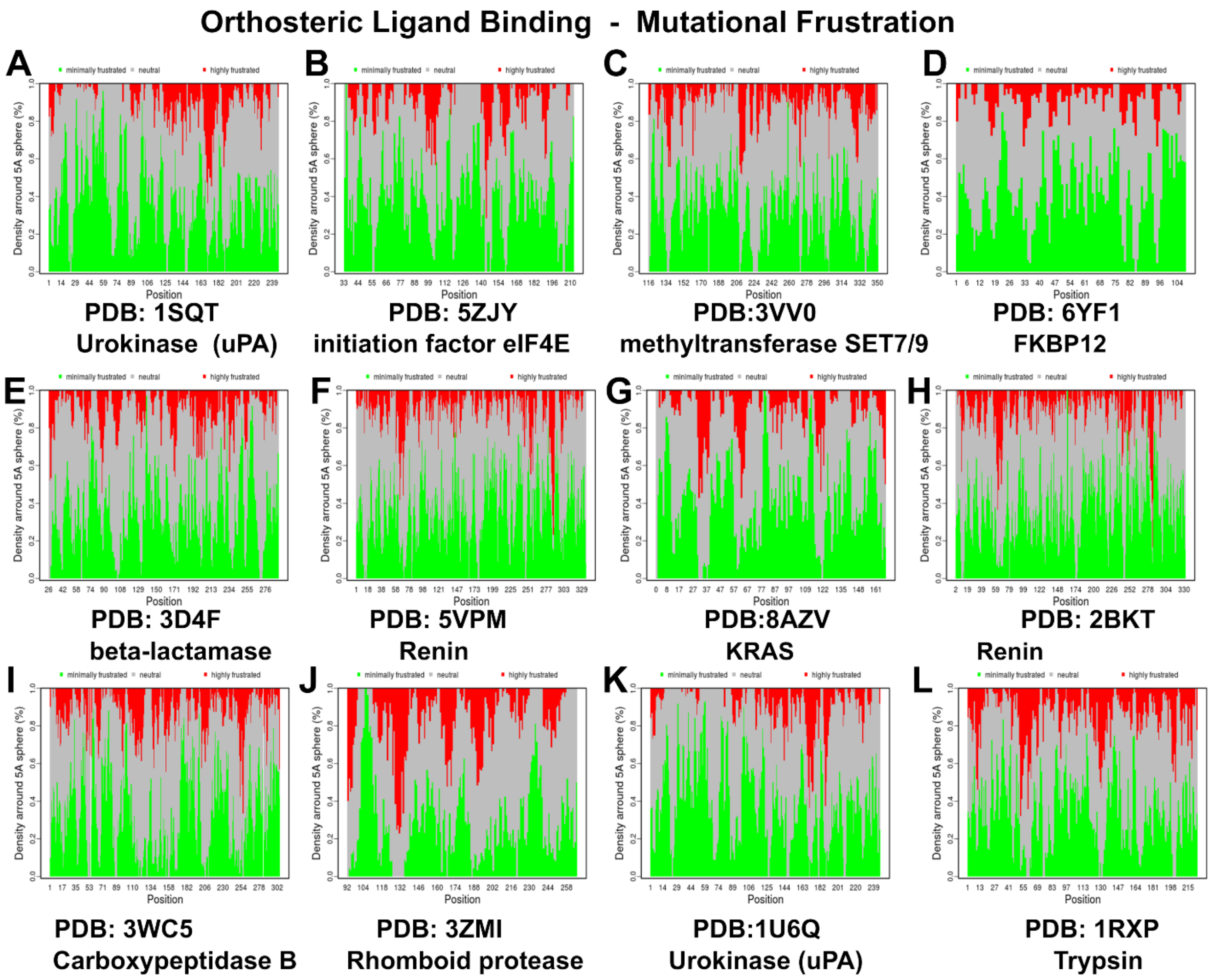


**Figure S6. Residue-level mutational frustration profiles for representative orthosteric ligand–protein complexes.** For each complex, the fraction of mutational frustration contact types is plotted per residue position: minimally frustrated (green), neutral (gray), and highly frustrated (red). Profiles are shown for (A) urokinase (uPA, PDB 1SQT), (B) initiation factor eIF4E (PDB 5ZJY), (C) methyltransferase SET7/9 (PDB 3VVO), (D) FKBP12 (PDB 6YF1), (E) beta-lactamase (PDB 3D4F), (F) renin (PDB 5VPM), (G) KRAS (PDB 8AZV), (H) renin (PDB 2BKT), (I) carboxypeptidase B (PDB 3WCI), (J) rhomboid protease (PDB 3ZMI), (K) urokinase (PDB 1U6Q), and (L) trypsin (PDB 1RXP). Residues comprising the orthosteric binding pocket consistently display elevated minimally frustrated fractions, indicating evolutionary constraint and optimized interaction energetics. The surrounding protein matrix exhibits predominantly neutral frustration, confirming that orthosteric sites are embedded within locally stabilized energetic environments.

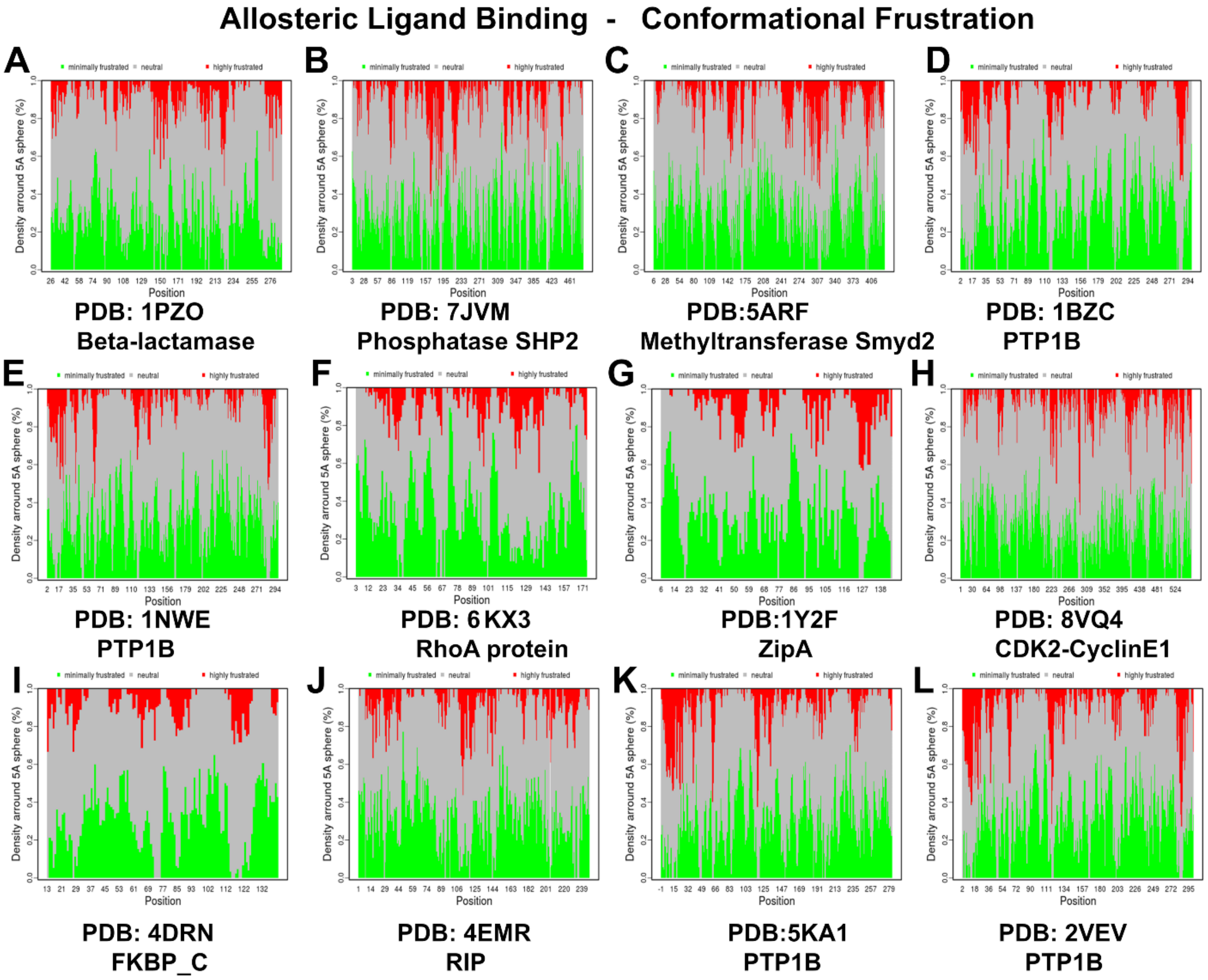


**Figure S7. Residue-level conformational frustration profiles for representative allosteric ligand–protein complexes.** For each complex, the fraction of mutational frustration contact types is plotted per residue position: minimally frustrated (green), neutral (gray), and highly frustrated (red). Profiles are shown are colored by residue-level conformational frustration state: minimally frustrated (green), neutral for a diverse set of complexes including (A) beta-lactamase (PDB 1PZO), (B) phosphatase SHP2 (PDB 7JVM), (C) methyltransferase Smyd2 (PDB 5ARF), (D) PTP1B (PDB 1BZC), (E) RhoA protein (PDB 6KX3), (G) ZipA (PDB 1Y2F), (H) CDK2-cyclin E1 (PDB 8VQ4), (I) FKBP_C (PDB 4DRN), (J) RIP (PDB 4EMR), (K) PTP1B (PDB 5KA1), and (L) PTP1B (PDB 2VEV). In contrast to orthosteric sites, allosteric binding pockets are characterized by pervasive neutral frustration with only sparse, non-contiguous minimally frustrated residues. The absence of concentrated minimal frustration networks reflects the conformational plasticity inherent to these regulatory regions.

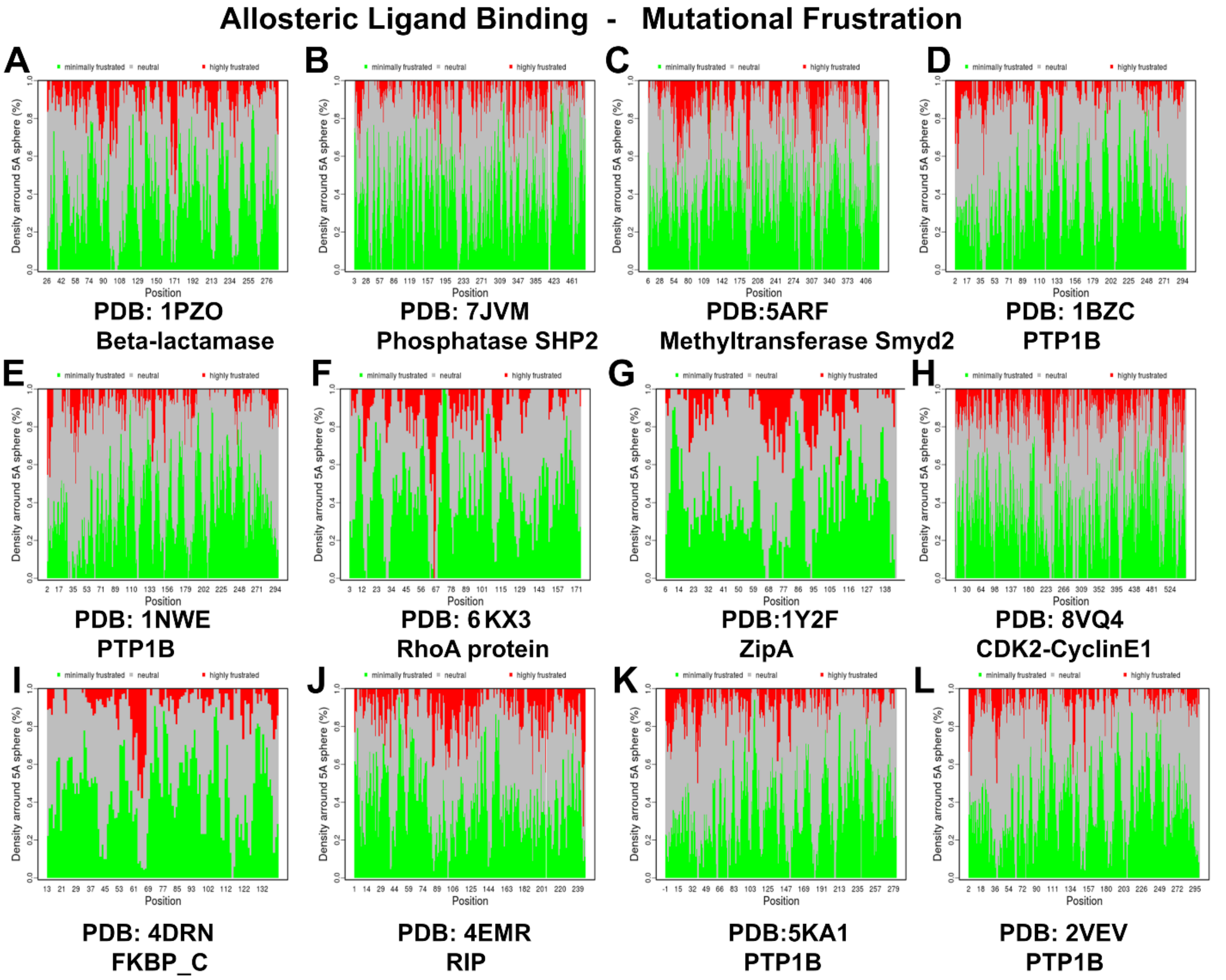


**Figure S7. Residue-level mutational frustration profiles for representative allosteric ligand–protein complexes.** For each complex, the fraction of mutational frustration contact types is plotted per residue position: minimally frustrated (green), neutral (gray), and highly frustrated (red). Profiles are shown are colored by residue-level conformational frustration state: minimally frustrated (green), neutral for a diverse set of complexes including (A) beta-lactamase (PDB 1PZO), (B) phosphatase SHP2 (PDB 7JVM), (C) methyltransferase Smyd2 (PDB 5ARF), (D) PTP1B (PDB 1BZC), (E) RhoA protein (PDB 6KX3), (G) ZipA (PDB 1Y2F), (H) CDK2-cyclin E1 (PDB 8VQ4), (I) FKBP_C (PDB 4DRN), (J) RIP (PDB 4EMR), (K) PTP1B (PDB 5KA1), and (L) PTP1B (PDB 2VEV). Allosteric pocket residues exhibit predominantly neutral frustration fractions across all positions, with only modest minimally frustrated contributions. This profile confirms evolutionary permissiveness and the absence of strong directional stabilization signals that pattern-based AI models require for accurate prediction.
